# Mechanism and regulation of the Bim1 interaction with the outer kinetochore Ndc80 complex in *S. cerevisiae*

**DOI:** 10.64898/2026.08.26.747248

**Authors:** Yvonne B. Winterborn, Christopher Batters, Tomos E. Morgan, Stefan M. V. Freund, David Barford

## Abstract

During eukaryotic cell division, kinetochores couple duplicated sister chromatids to mitotic spindle microtubules to mediate faithful chromosome segregation. Although the main kinetochore attachment sites to centromeric chromatin and microtubules are known, additional factors including microtubule-associated proteins are required for efficient chromosome biorientation and segregation *in vivo*. However, the roles and mechanisms of these factors in kinetochore function remain to be fully understood. Here, we characterise a previously unrecognised interaction between the microtubule plus-end tracking protein Bim1 and the outer kinetochore Ndc80 complex (Ndc80c) in *S. cerevisiae*. We show this interaction is mediated by a conserved SxIP motif within the intrinsically disordered Ndc80 N-terminus (Ndc80^N^), augmented by a secondary binding site containing an α-helical segment. This Ndc80 interaction with Bim1 increases the strength of Ndc80c-microtubule attachments. Phosphorylation of the Bim1-binding region of Ndc80^N^ by the error correction Ipl1/Aurora B protein kinase alters its secondary structure and weakens the Bim1–Ndc80c interaction, providing a potential additional regulatory mechanism for how incorrect kinetochore-microtubule attachments are destabilised during error correction.

## Introduction

Kinetochores are large multiprotein complexes that assemble at centromeres and bind spindle microtubules, forming load-bearing attachments between chromosomes and the mitotic spindle, thereby ensuring accurate chromosome segregation (McAinsh & Marston, 2022; Musacchio & Desai, 2017). The inner kinetochore constitutive centromere-associated network (CCAN) complex assembles at centromere-specific CENP-A nucleosomes, topologically entrapping DNA (Dendooven et al., 2023; Yatskevich et al., 2022). CCAN recruits the outer kinetochore KMN network complex, comprised of the Knl1 (Knl1c), Mtw1 (Mtw1c) and Ndc80 (Ndc80c) sub-complexes (Cheeseman et al., 2006). Knl1c functions as a signalling scaffold for the spindle assembly checkpoint (SAC) which coordinates anaphase onset with the correct attachment of all chromosomes to the spindle (Fischer, 2023; McAinsh & Kops, 2023). Mtw1c organises the KMN complex and its recruitment to kinetochores (Petrovic et al., 2010; Polley et al., 2024; Turner et al., 2026; Yatskevich et al., 2024). Ndc80c is the primary microtubule-binding component of KMN (Tooley & Stukenberg, 2011; Wei et al., 2007). Kinetochore-microtubule coupling is augmented by the outer kinetochore Dam1 complex (Dam1c) in budding yeast and Ska and Astrin-SKAP complexes in vertebrates (Helgeson et al., 2018; Huis in ’t Veld et al., 2019; Kern et al., 2017; Lampert et al., 2010; Muir et al., 2023; Tien et al., 2010).

Ndc80c comprises Ndc80, Nuf2, Spc24 and Spc25 (Wigge & Kilmartin, 2001). The complex binds to microtubules through the N-terminal calponin homology (CH) domains of Ndc80 and Nuf2 (Alushin et al., 2010; Muir et al., 2023), whereas Spc24 and Spc25 connect Ndc80c to Mtw1c within the KMN complex (Polley et al., 2024; Turner et al., 2026; Wei et al., 2006; Yatskevich et al., 2024). The N-terminal 114 residues of Ndc80 (Ndc80^N^) are intrinsically disordered and contribute to Ndc80c microtubule binding (Suzuki et al., 2016; Wei et al., 2007). Dam1c is a heterodecameric complex that oligomerises into rings of 16 complexes that encircle microtubules (Cheeseman et al., 2001; Miranda et al., 2005; Muir et al., 2023; Ramey et al., 2011; Umbreit et al., 2014; Westermann et al., 2005). The three interaction sites between Ndc80c and Dam1c (Kim et al., 2017; Muir et al., 2023; Zahm et al., 2023) augment Ndc80c-microtubule attachment strength (Muir et al., 2023; Tien et al., 2010; Umbreit et al., 2014), whereas the Dam1c ring enables Ndc80c to track depolymerising microtubule plus-ends (Lampert et al., 2010; Tien et al., 2010).

Ndc80c and Dam1c are the major targets of the error correction mechanism, whose effector is the Ipl1/Aurora B protein kinase. This mechanism functions to reset kinetochore-microtubule attachments that fail to establish bioriented attachment to the mitotic spindle (Biggins & Murray, 2001; Cheeseman et al., 2002; Doodhi et al., 2021; Pinsky et al., 2006; Tanaka et al., 2002). Phosphorylation of Dam1c by Ipl1 weakens Ndc80c-Dam1c interactions and suppresses Dam1c ring formation (Cheeseman et al., 2002; Kim et al., 2017; Muir et al., 2023; Tien et al., 2010). Ndc80^N^ also contains multiple Ipl1 phosphorylation sites. Phosphorylation at these sites weakens Ndc80c microtubule binding, likely because the negatively charged phosphate groups reduce the affinity of the positively charged Ndc80^N^ for the negatively charged tubulin tails (Akiyoshi et al., 2009; Cheeseman et al., 2002; Doodhi et al., 2021; Wei et al., 2007). Upon biorientation, kinetochores come under tension, promoting dephosphorylation of the outer kinetochore and stabilisation of kinetochore-microtubule attachments, thereby satisfying the SAC for progression into anaphase (Lara-Gonzalez et al., 2021; Pinsky & Biggins, 2005).

Although the main kinetochore components and microtubule binding sites are defined, additional factors are required to ensure accurate and efficient chromosome segregation *in vivo*, reviewed in (Joglekar & Kukreja, 2017; McAinsh & Marston, 2022). Many of these factors are microtubule-associated proteins (MAPs), which play roles in chromosome capture, establishment of end-on attachments and chromosome biorientation (Tanaka, 2010). Direct MAP-kinetochore interactions have previously been described for Stu2/ch-TOG with Ndc80c, and for Bim1/EB1 with the Dam1, Ska and Astrin/SKAP complexes (Dudziak et al., 2021; Dunsch et al., 2011; Miller et al., 2016; Miller et al., 2019; Radhakrishnan et al., 2023; Thomas et al., 2016). The budding yeast Bim1 (binding to microtubules) is an autonomous plus-end tracking protein (+TIP) of the end-binding (EB) protein family. Bim1 localises to growing microtubule plus-ends, and functions as a master regulator of the +TIP network, recruiting a range of binding partners to microtubule plus-ends to regulate microtubule function and dynamics (Akhmanova & Steinmetz, 2008; Blake-Hodek et al., 2010; Schwartz et al., 1997; Stangier et al., 2018; Zimniak et al., 2009). Bim1 aids Dam1c localisation during the formation of end-on attachments and promotes Dam1c ring formation (Dudziak et al., 2021).

In this study, we combined NMR, biochemical and biophysical approaches with protein structure prediction to mechanistically and structurally characterise a previously unrecognised interaction between Bim1 and Ndc80c. We show this interaction is mediated by recognition of a conserved SxIP motif within Ndc80^N^ by the Bim1 end-binding homology (EBH) domain, while a distinct secondary binding site within Ndc80^N^ contributes to the interaction. Bim1 increases the strength of Ndc80c-microtubule attachments and confers plus-end tracking properties on Ndc80c. Furthermore, the Bim1–Ndc80c interaction is regulated by Ipl1, providing potential insight into how incorrect kinetochore-microtubule attachments may be destabilised during error correction.

## Results

### Bim1 interacts with the outer kinetochore Ndc80 complex

We purified full-length Bim1 and Ndc80c from insect cells and tested their interaction *in vitro* by analytical size-exclusion chromatography (SEC). When mixed at an equimolar ratio of 10 µM, Bim1 showed a clear shift in elution volume to co-elute with Ndc80c, indicating complex formation (Figure 1A). The concentrations of Bim1 and Bim1 variants used in this study are stated as a dimer. Both the chromatogram and SDS-PAGE gels revealed excess unbound Bim1, suggesting either that the Bim1:Ndc80c complex does not form with strict 1:1 stoichiometry or that the complex partially dissociates under these conditions. To distinguish between these possibilities, we analysed the complex by SEC-MALS. Although some dissociation was observed, the measured molecular mass of the complex was 254 kDa (Figure 1B), consistent with formation of a 1:1 complex between a Bim1 dimer (76 kDa) and Ndc80c (183 kDa). The low-affinity interaction between Ndc80c and Bim1 may explain the absence of complex formation in a previous study (Dudziak et al., 2021).

**Figure 1.**
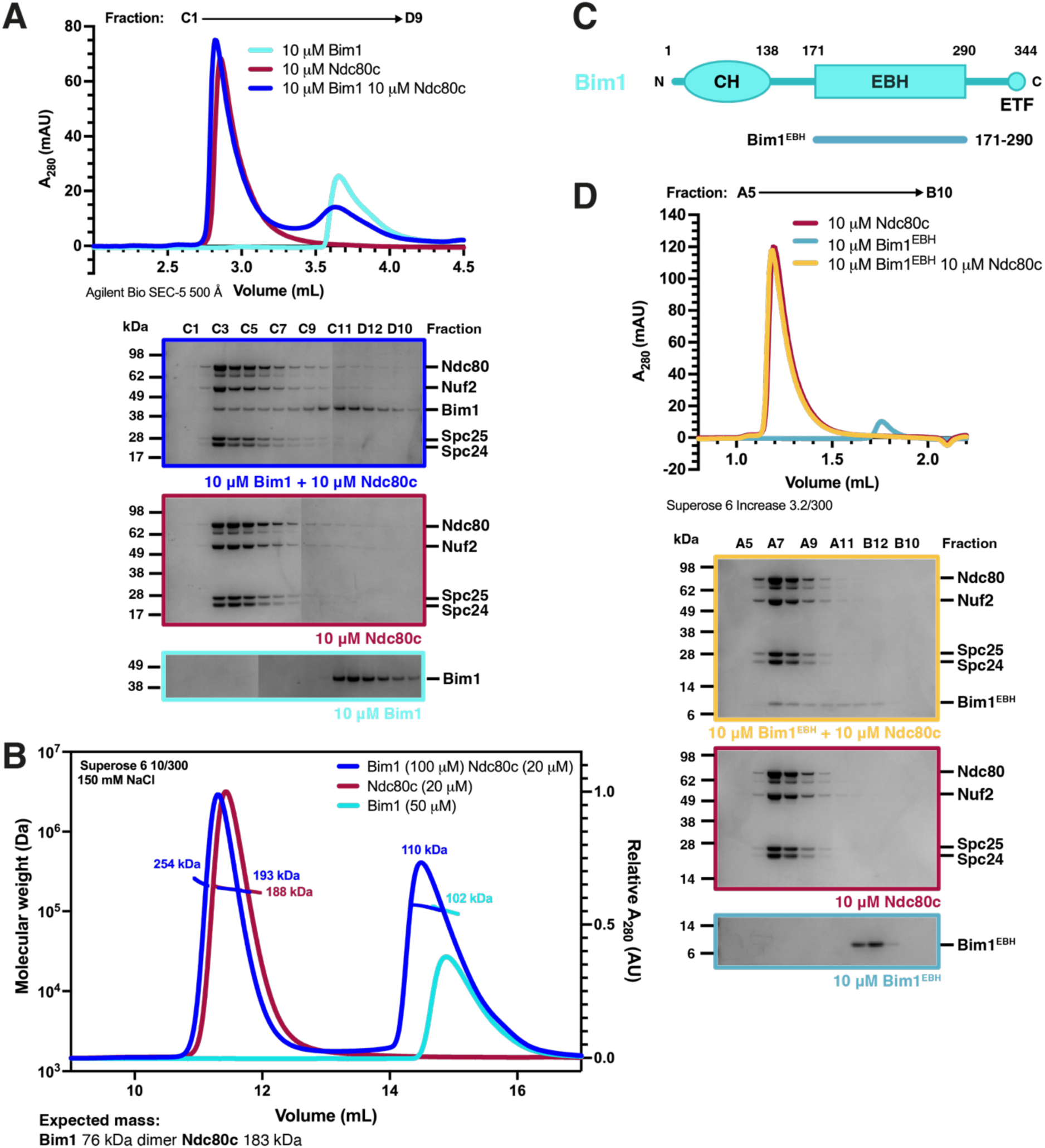
– Bim1 binds to the Ndc80c, mediated by the Bim1 EBH domain. **(A)** Chromatogram (top) and corresponding SDS-PAGE analysis (bottom) of eluate fractions of Bim1 and Ndc80c injected onto an Agilent Bio SEC-5 500 Å column individually and together. **(B)** SEC-MALS analysis of the Bim1–Ndc80c interaction. The Bim1:Ndc80c trace indicates some dissociation of the complex in these conditions; the front of the peak gives an Mr of 254 kDa, corresponding to one Ndc80c interacting with one Bim1 dimer, and the tail of the peak gives an Mr of 193 kDa, corresponding to Ndc80c alone. **(C)** Domain architecture of Bim1. The Bim1^EBH^ is comprised of residues 171-290 of Bim1. CH = calponin homology, EBH = end-binding homology, ETF = glutamic acid-threonine-phenylalanine. **(D)** Chromatogram (top) and corresponding SDS-PAGE analysis (bottom) of eluate fractions of Bim1^EBH^ and Ndc80c injected onto a Superose 6 Increase 3.2/300 column individually and together at equimolar concentrations.

Bim1 interacts with many of its binding partners, including Dam1c (Dudziak et al., 2021), through its central EBH domain, which also mediates its dimerisation (Figure 1C). We therefore purified the Bim1 EBH domain (Bim1^EBH^: residues 171-290) and tested its interaction with Ndc80c by analytical SEC at an equimolar ratio of 10 µM. As with full-length Bim1, Bim1^EBH^ shifted to co-elute with Ndc80c, indicating that the EBH domain is sufficient to mediate the interaction (Figure 1D).

### A conserved SxIP motif in Ndc80^N^ recognises Bim1^EBH^

EBH domains recognise SxIP and LxxPTPh (h: hydrophobic) motifs (Honnappa et al., 2009; Kumar et al., 2017). Multiple sequence alignment of budding yeast Ndc80 identified a conserved SQIP motif in Ndc80^N^ (residues 22-25 of *S. cerevisiae* Ndc80) (Figure 2A). Deletion of Ndc80^N^ (residues 1-114) or the Ndc80^N^ SQIP motif from Ndc80c (Ndc80c^Ndc80-ΔN^ and Ndc80c^ΔSQIP^, respectively) abolished the interactions between Ndc80c and Bim1, as assessed by analytical SEC (Figure 2B and Figure S1A). These results suggested that the Bim1–Ndc80c interaction is mediated through a canonical EBH-SxIP recognition mechanism involving Bim1^EBH^ and the Ndc80^N^ SQIP motif.

**Figure 2.**
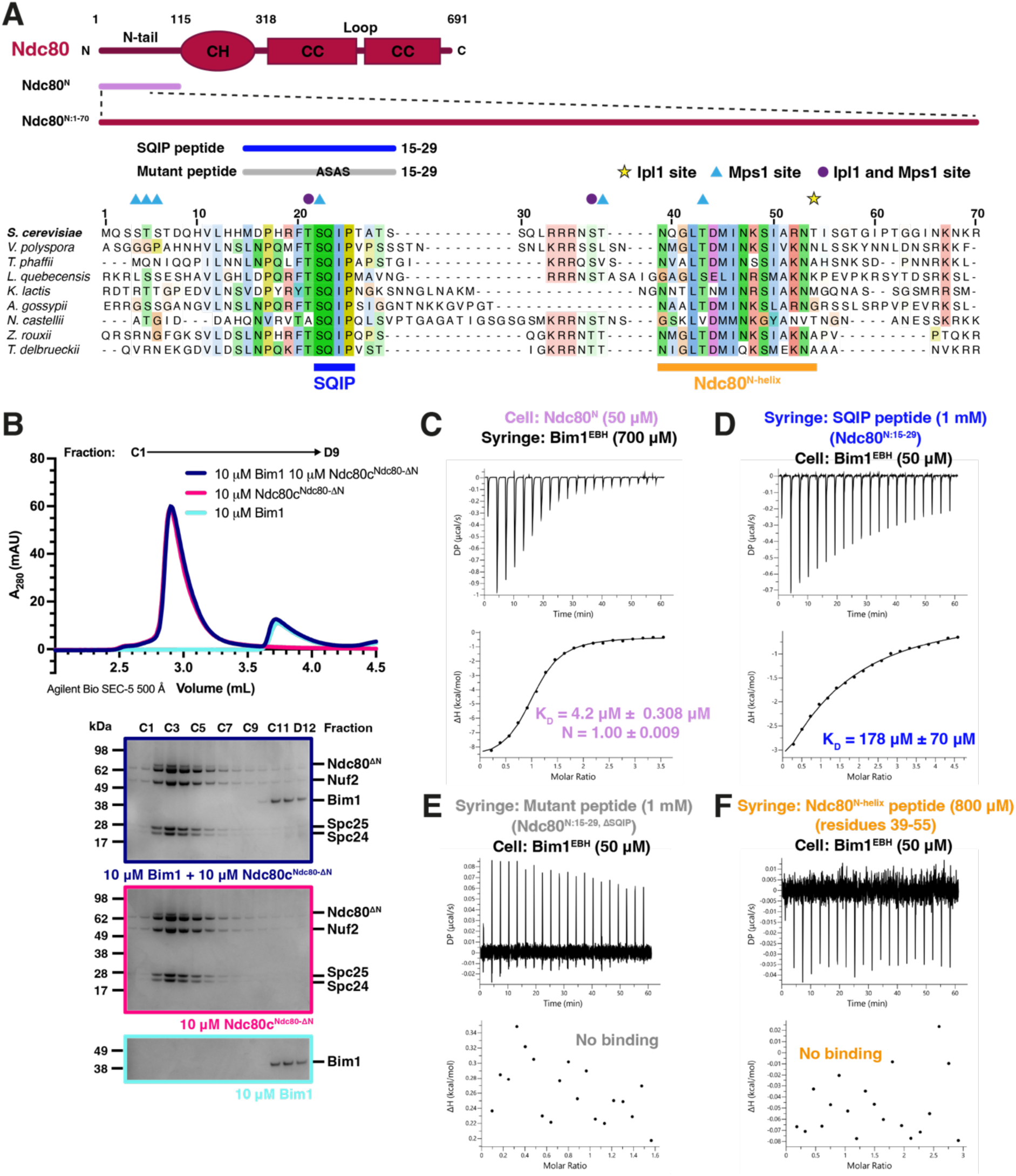
– A conserved SxIP motif in Ndc80^N^ drives the Bim1-Ndc80c interaction. **(A)** Domain architecture of Ndc80 and variants used in this study (top). Ndc80^N^ comprises the entire disordered Ndc80^N^, Ndc80^N:1-70^ contains the full Bim1-binding region of the Ndc80^N^. SQIP and mutant peptides comprise Ndc80 residues 15-29, the mutant peptide has residues 22-25 (SQIP) replaced with ASAS. Multiple sequence alignment across budding yeast of Ndc80^N:1-70^ (bottom), with Ipl1 and Mps1 phosphorylation sites indicated. Conserved regions include the SQIP motif and a predicted helix (Ndc80^N-helix^). CH: calponin homology, CC: coiled-coil. **(B)** Chromatogram (top) and corresponding SDS-PAGE analysis (bottom) of eluate fractions of Bim1 and Ndc80c^Ndc80-ΔN^ (Ndc80^N^ deleted from Ndc80c) injected onto an Agilent Bio SEC-5 500 Å column individually and together at equimolar concentrations. **(C)** ITC of Ndc80^N^ with Bim1^EBH^. KD: dissociation constant, N: stoichiometry. **(D)** ITC of the SQIP with Bim1^EBH^. Stoichiometry could not be determined due to a low c-value. **(E)** ITC of the mutant peptide with Bim1^EBH^. **(F)** ITC of the Ndc80^N-helix^ peptide, residues 39-55 of Ndc80^N^, with Bim1^EBH^.

To quantify the affinity of the Bim1–Ndc80c interaction, we performed isothermal titration calorimetry (ITC) using Bim1^EBH^ and Ndc80^N^. This revealed an interaction affinity (K_D_) of 4.2 µM and a stoichiometry of one Ndc80^N^ per Bim1^EBH^ dimer (Figure 2C and Table S1), consistent with the stoichiometry of the full complex determined using SEC-MALS and with the partial dissociation of the Bim1:Ndc80c complex in our SEC experiments (Figure 1A). This affinity is comparable to other known SxIP-mediated interactions, including the human EB1:MCAF and EB1:APC complexes (Almeida et al., 2024; Honnappa et al., 2009). +TIPs such as Bim1 are concentrated at microtubule plus-ends, and their micromolar interaction affinities are thought to permit dynamic remodelling of the +TIP network (Akhmanova & Steinmetz, 2008, 2015).

For some +TIP complexes, the SxIP motif alone is sufficient to confer full interaction affinity, whereas in others, additional residues are required (Almeida et al., 2024; Buey et al., 2012; Honnappa et al., 2005), reviewed in (Kumar & Wittmann, 2012). Using ITC, we found that a synthetic SQIP peptide corresponding to Ndc80 residues 15-29 (Figure 2A) bound Bim1^EBH^ with a K_D_ of 178 µM, with one SQIP peptide binding per Bim1^EBH^ dimer (Figure 2D and Table S1). In contrast, a peptide containing a mutated SQIP motif showed no detectable binding to Bim1^EBH^ by ITC (Figure 2E and Table S1). These results demonstrate that the SQIP motif is important for the Bim1–Ndc80c interaction, but also indicate that additional residues within Ndc80^N^ are required to achieve the low micromolar-affinity interaction. To investigate the molecular basis of the Bim1–Ndc80c interaction, we combined protein structure prediction with NMR spectroscopy.

### Structure prediction of the Bim1^EBH^-Ndc80^N^ interaction

We used AlphaFold2 (Jumper et al., 2022) to predict the structure of an *S. cerevisiae* Bim1^EBH^ dimer:Ndc80^N^ complex at 1:1 stoichiometry (Figure 3). Bim1^EBH^ was predicted with high confidence and low predicted alignment errors (PAEs) and, consistent with the crystal structure of the Bim1 EBH domain (Hüls et al., 2012), adopts a coiled-coil architecture that forms a four-helical bundle at the C-terminus of the domain (Figure 3A, D). The Ndc80 SQIP motif was also predicted with high confidence (pLDDT > 90) and low PAE (< 5 Å) to bind one of the SxIP-binding sites of the Bim1^EBH^ dimer (Figure 3A, B), consistent with experimentally determined SxIP:EBH structures (Honnappa et al., 2009; Kumar et al., 2017). Although the remainder of Ndc80^N^ was predicted with lower confidence, the model suggested that Ndc80^N^ wraps around Bim1^EBH^ and positions an α-helix (residues 39-54) at the second SxIP binding site of the Bim1^EBH^ dimer, with a PAE of 5-10 Å (Figure 3A, C, D).

**Figure 3.**
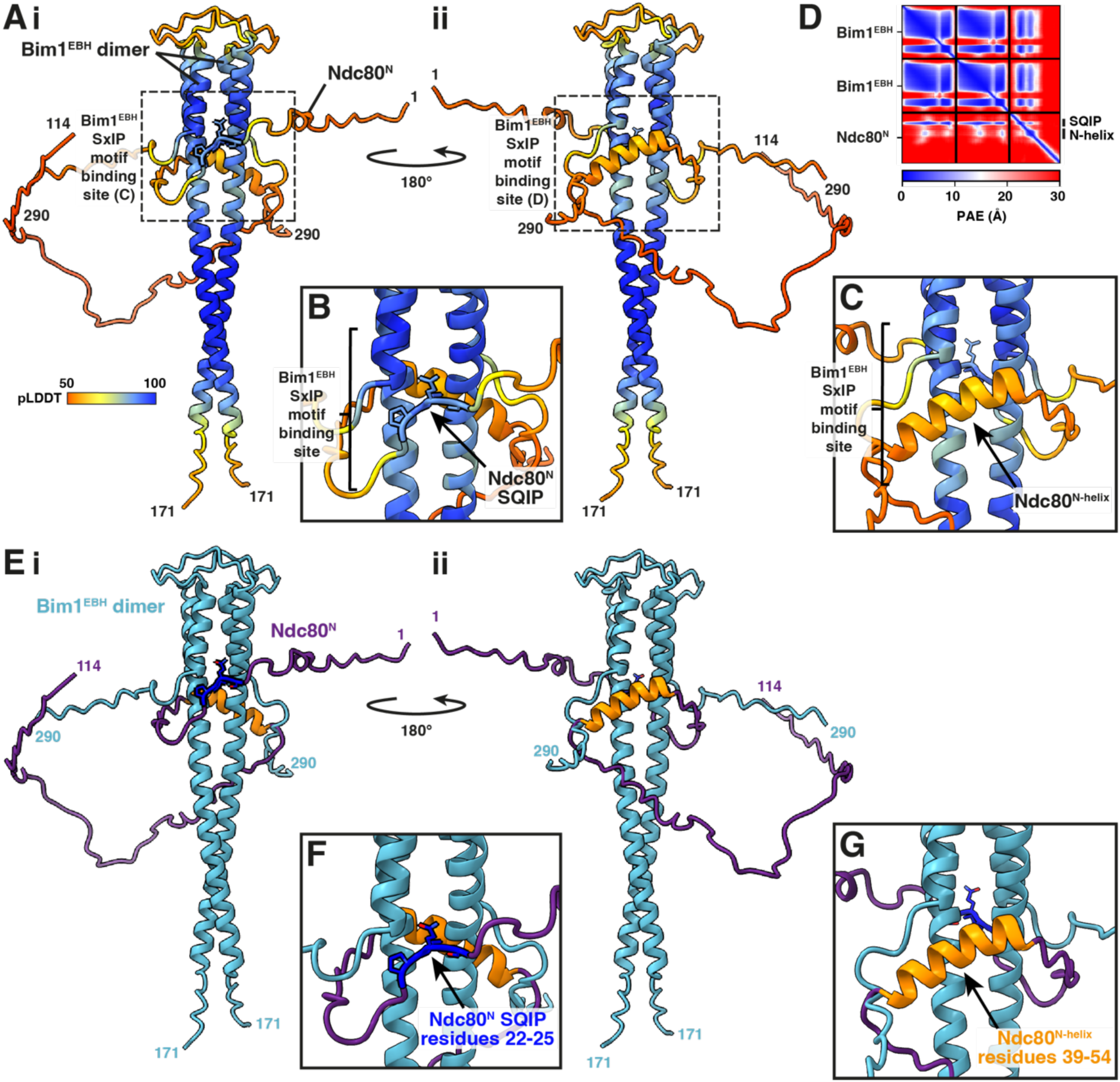
– Structure prediction of the Bim1^EBH^-Ndc80^N^ interaction. **(A)** Bim1^EBH^ dimer and Ndc80^N^ AlphaFold2 prediction coloured by pLDDT, viewing the two SxIP motif binding sites of the Bim1^EBH^ dimer (i and ii). **(B)** Close-up from A(i) of the Ndc80^N^ SQIP motif predicted with high confidence to bind one SxIP motif binding site of Bim1^EBH^. **(C)** Close-up from A(ii) of the predicted positioning of the low confidence Ndc80^N-helix^ close to the other SxIP motif binding site of Bim1^EBH^. **(D)** PAE plot for the Bim1^EBH^-Ndc80^N^ prediction. **(E-G)** Bim1^EBH^ dimer and Ndc80^N^ AlphaFold2 prediction and close-ups as in (A-C), coloured by chain.

The residues in the predicted helix (Ndc80^N-helix^) lie within a conserved region of Ndc80^N^ (Figure 2A). ITC measurements using a peptide corresponding to Ndc80^N-helix^ failed to detect binding to Bim1^EBH^ (Figure 2F), however the comparable degree of sequence conservation of Ndc80^N-helix^ and the SQIP motif supports the idea that Ndc80^N-helix^ contributes to Bim1–Ndc80c interactions. To test the AlphaFold model, we next used NMR spectroscopy to obtain mechanistic insight into the residue-specific interactions mediating the Bim1^EBH^:Ndc80c complex.

### Optimisation of a system to investigate the Bim1–Ndc80c interaction by NMR

Titration of unlabelled Bim1^EBH^ into ^15^N-labelled Ndc80^N^ showed that the Bim1-binding region of Ndc80^N^ spans residues 10-64 (Figure S2A-C). However, the Ndc80^N^ backbone assignment within this region remained incomplete due to low signal intensity and spectral overlap. In addition, the elongated coiled-coil structure of Bim1^EBH^ makes it highly anisotropic and slow-tumbling in solution, inducing rapid transverse relaxation (T_2_) and resulting in poor resonance sensitivity. Consequently, many Bim1^EBH^ resonances remained unassigned even with the enhanced sensitivity afforded by fractional side-chain deuteration (Figure S3D). Binding to an anisotropic unlabelled partner also causes substantial line broadening, complicating interpretation of the extent to which individual residues contribute to the interaction (Figure S2C). To improve spectral quality, we generated a truncated Ndc80^N^ (residues 1-70: Ndc80^N:1-70^). Ndc80^N:1-70^ bound Bim1^EBH^ with an affinity of 4.0 µM, identical to full-length Ndc80^N^, and, as predicted by AlphaFold2, formed equivalent interactions with Bim1^EBH^ (Figure 2A, Figure S2E, F and Table S1). In the 2D ^1^H,^15^N heteronuclear single quantum coherence (HSQC) spectrum of Ndc80^N:1-70^, all backbone amide resonances except those corresponding to Ser29, Arg35, and proline residues were visible and assigned using standard triple resonance methods (Figure 4A). We also designed a shortened Bim1^EBH^ protein, Bim1^EBH-mini^ (residues 205-285), which retains the complete SxIP-binding site while containing a truncated coiled-coil region (Figure S3B). Bim1^EBH-mini^ bound with similar affinity as Bim1^EBH^ to Ndc80^N:1-70^ (Figure S3C and Table S1). The improved relaxation behaviour of Bim1^EBH-mini^ enabled a complete backbone resonance assignment (Figure S3D). ^15^N transverse relaxation (T_2_) experiments, analysed as R_2_ rates (1/T_2_), showed that the less dynamic regions of Bim1^EBH-mini^ correspond to the predicted coiled-coil segments (Figure S3E), confirming that Bim1^EBH-mini^ adopts the same coiled-coil architecture as the corresponding region of Bim1^EBH^.

**Figure 4.**
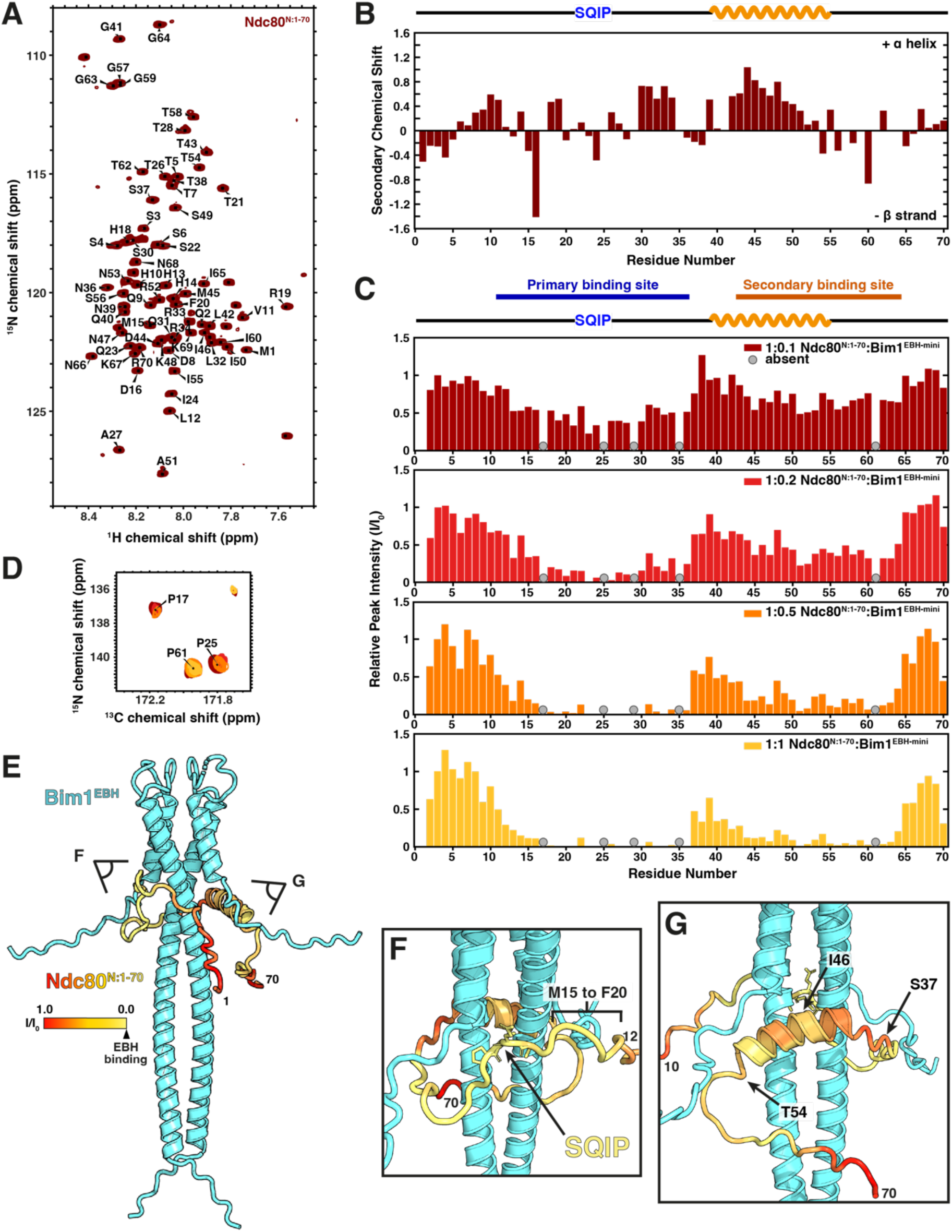
– NMR titration experiments reveal primary and secondary Bim1-binding sites in the Ndc80 N-tail. **(A)** 2D ^1^H, ^15^N HSQC spectra of ^15^N-labelled Ndc80^N:1-70^ with backbone assignments determined using a ^13^C,^15^N-labelled sample. All amide resonances could be assigned except Ser29 and Arg35. **(B)** Secondary structure propensity based on calculated secondary Cα, Cβ chemical shifts of ^13^C,^15^N-labelled Ndc80^N:1-70^. Positive values indicate α-helical propensity and negative values indicate β-strand propensity. Regions of Ndc80^N:1-70^, including the predicted Ndc80^N-helix^, sample α-helical secondary structure. Large negative shifts for Asp16, Ile24 and Ile60 are the consequence of preceding prolines. **(C)** Relative peak intensities (I/I0) of peaks from the ^1^H, ^15^N 2D HSQC spectra of Ndc80^N:1-70^ titrated with the indicated concentrations of Bim1^EBH-mini^. Lower relative peak intensity in the presence of Bim1^EBH-mini^ is due to peak attenuation, indicating residues involved in binding. A primary binding site comprises the SQIP motif and surrounding residues (residues 11-36), and a secondary binding site comprises Ndc80^N-helix^ and additional residues C-terminal to it (residues 42-64). ‘Absent’ indicates peaks which are not assigned in the ^1^H, ^15^N 2D HSQC spectra of free Ndc80^N:1-70^, including three prolines which lack amide resonances. Intensities are normalised relative to Ser3. **(D)** Overlay of the proline section of the ^13^C, ^15^N 2D CON spectra of ^13^C,^15^N-labelled Ndc80^N:1-70^ titrated with Bim1^EBH-mini^ as in (A), showing the peak attenuation observed for Pro17 and Pro25, and to a lesser extent for Pro61. **(E)** Relative peak intensities (I/I0) of 1:0.5 Ndc80^N:1-70^:Bim1^EBH-mini^ plotted onto the AlphaFold2 prediction of the Ndc80^N:1-70^-Bim1^EBH^ interaction. Ndc80^N:1-70^ is coloured by I/I0. **(F)** Close-up of the SQIP motif from (E). Low I/I0 values for Met15-Phe20 suggest they are positioned closer to Bim1^EBH^ in the interaction than predicted. **(G)** Close-up of the predicted secondary binding site from (E). NMR titration binding data suggest that the centre of the secondary interaction site is around Thr54, not Ile46 as predicted. The true position of Ndc80^N^ in the Bim1-Ndc80c interaction may be slightly shifted compared to the AlphaFold2 prediction.

### Ndc80^N^ engages the SxIP motif-binding site of Bim1

2D ^1^H,^15^N HSQC-based titration of unlabelled Ndc80^N^ into ^15^N-labelled Bim1^EBH-mini^ caused substantial peak attenuation for amide resonances of residues located within and surrounding the SxIP motif-binding site (Figure S3F). Peak attenuation in the presence of an unlabelled binding partner indicates residues that participate in, or are affected by, the binding interaction. This is a result of exchange between a free and bound state (for micromolar-affinity binding) and increased transverse relaxation due to increased molecular weight of the bound state. Attenuation was quantified as the relative peak intensity (I/I_0_) and mapped onto the AlphaFold2 model of the Bim1^EBH-mini^:Ndc80^N^ complex (Figure S3F). Residues showing the greatest attenuation clustered within and around the SxIP-binding site, in strong agreement with the AlphaFold2 prediction.

### Ndc80^N^ contains primary and secondary Bim1-binding sites

NMR spectroscopy was used to investigate the structural and dynamic properties of Ndc80^N:1-70^ and its interaction with Bim1^EBH-mini^. The narrow dispersion of ^1^H chemical shifts in the 2D ^1^H,^15^N HSQC spectrum of Ndc80^N:1-70^ (Figure 4A) confirmed that it is largely disordered in solution (Okazaki et al., 2018). To assess the α-helical propensity of the predicted Ndc80^N-helix^, we analysed residue-specific secondary chemical shifts based on the assignment of Cα and Cβ resonances for each residue. These provide a reliable measure of secondary structure propensity in disordered proteins (Spera & Bax, 1991). Residues corresponding to the predicted Ndc80^N-helix^ exhibited positive secondary chemical shifts, consistent with α-helical propensity (Figure 4B). Additionally, residues 6-11 and 30-34 showed weak α-helical propensity.

Titration of unlabelled Bim1^EBH-mini^ into ^15^N-labelled Ndc80^N:1-70^ caused substantial peak attenuation for amide resonances in the 2D ^1^H,^15^N HSQC spectra (Figure S4A). Peak attenuation was quantified as the relative peak intensity (I/I_0_), normalised to Ser3, a residue unaffected by Bim1^EBH-mini^ binding (Figure 4C). These experiments identified two distinct regions of Ndc80^N^ that interact with Bim1. The strongest attenuation was centred on the SQIP motif and its surrounding residues (residues 15-36). A second region of attenuation was detected within and immediately C-terminal to the Ndc80^N^ helix (residues 42-64) (Figure 4C). We therefore designated these regions as the primary and secondary Bim1-binding sites, respectively (Figure 4C). Peak attenuation within the primary binding site was observed at low concentrations of Bim1^EBH-mini^, progressing to an almost complete loss of signal at an Ndc80^N:1-70^:Bim1^EBH-mini^ ratio of 1:0.5 (Figure 4C). In contrast, residues within the secondary binding site showed weaker attenuation at low Bim1^EBH-mini^ concentrations (Figure 4C). The two binding sites are separated by residues 37-41, which were only partially attenuated even in a stoichiometric Bim1^EBH-mini^:Ndc80^N:1-70^ complex. Carbon-detect titration experiments further supported these observations. Signals corresponding to Pro17 and Pro25, both located within the primary binding site (with Pro25 forming the proline of the SQIP motif), were strongly attenuated, whereas Pro61, located within the secondary binding site, showed weaker attenuation (Figure 4D), consistent with the results obtained for amide resonances of surrounding residues. Additionally, crosslinking mass spectrometry (CL-MS) of a Bim1^EBH^:Ndc80^N:1-70^ complex showed many crosslinks between residues within or close to the proposed primary and secondary Bim1-binding sites of Ndc80^N:1-70^ and the Bim1^EBH^ SxIP binding site (Figure S4B, C).

Mapping the I/I_0_ values obtained at a 1:0.5 ratio of Ndc80^N:1-70^ to Bim1^EBH-mini^ onto our AlphaFold2 model showed that residues of Ndc80^N:1-70^ with the lowest I/I_0_ values are located closest to Bim1^EBH^ (Figure 4E-G). The I/I_0_ values for the SQIP motif and its surrounding residues correlated well with the predicted model (Figure 4F). Residues Met15-Phe20 of the primary binding site immediately N-terminal to the SQIP motif also displayed low I/I_0_ values, suggesting that they are positioned closer to Bim1^EBH^ than predicted by AlphaFold2 (Figure 4F). This discrepancy is consistent with the low confidence scores and elevated PAEs for this region. The mapped I/I_0_ values also indicated that the secondary binding site is shifted C-terminal relative to the AlphaFold2 prediction. In the model, the centre of the secondary binding site is positioned at Ile46, near the middle of the Ndc80^N^ helix, and structurally equivalent to Ile of the SQIP motif (Figure 4G). However, our titration data identified Thr54, at the C-terminus of Ndc80^N-helix^, as the centre of the secondary binding site (Figure 4C). Given the low confidence and high PAE scores in this region, these inaccuracies in the AlphaFold2 prediction are not unexpected (Figure 3A, C, D).

To investigate whether Ndc80^N-helix^ becomes stabilised upon complex formation, we calculated the secondary chemical shifts of double-labelled Ndc80^N:1-70^ in the presence of Bim1^EBH-mini^ at molar ratios of 1:0.2, 1:0.5, and 1:1 (Figure S4D). At ratios above 1:0.2, many resonances were absent due to peak attenuation in the required 3D triple resonance experiments. Comparison of the secondary structure propensities showed no detectable changes upon binding Bim1^EBH-mini^ (Figure 4B and Figure S4D), indicating that the interaction with Bim1 does not alter the secondary structure of Ndc80^N:1–70^.

### The primary Bim1-binding site dominates Bim1–Ndc80c interaction affinity

Having defined the primary and secondary binding sites for Bim1^EBH^ within Ndc80^N^, we next used ITC to determine their relative contributions to the affinity of the Ndc80^N^-Bim1^EBH^ interaction. A peptide incorporating the primary binding site (residues 1-39) bound Bim1^EBH^ with a K_D_ of 21.5 µM (Figure 5A and Table S1), an eight-fold higher affinity than the SQIP peptide (residues 15-29), but a five-fold lower affinity than Ndc80^N:1-70^. In contrast, we were unable to determine a dissociation constant for a peptide corresponding to the secondary binding site (residues 40-70) (Figure 5B). These results indicate that, although the secondary binding site contributes to the interaction between Bim1^EBH^ and Ndc80^N^, it has a low intrinsic affinity for Bim1^EBH^.

**Figure 5.**
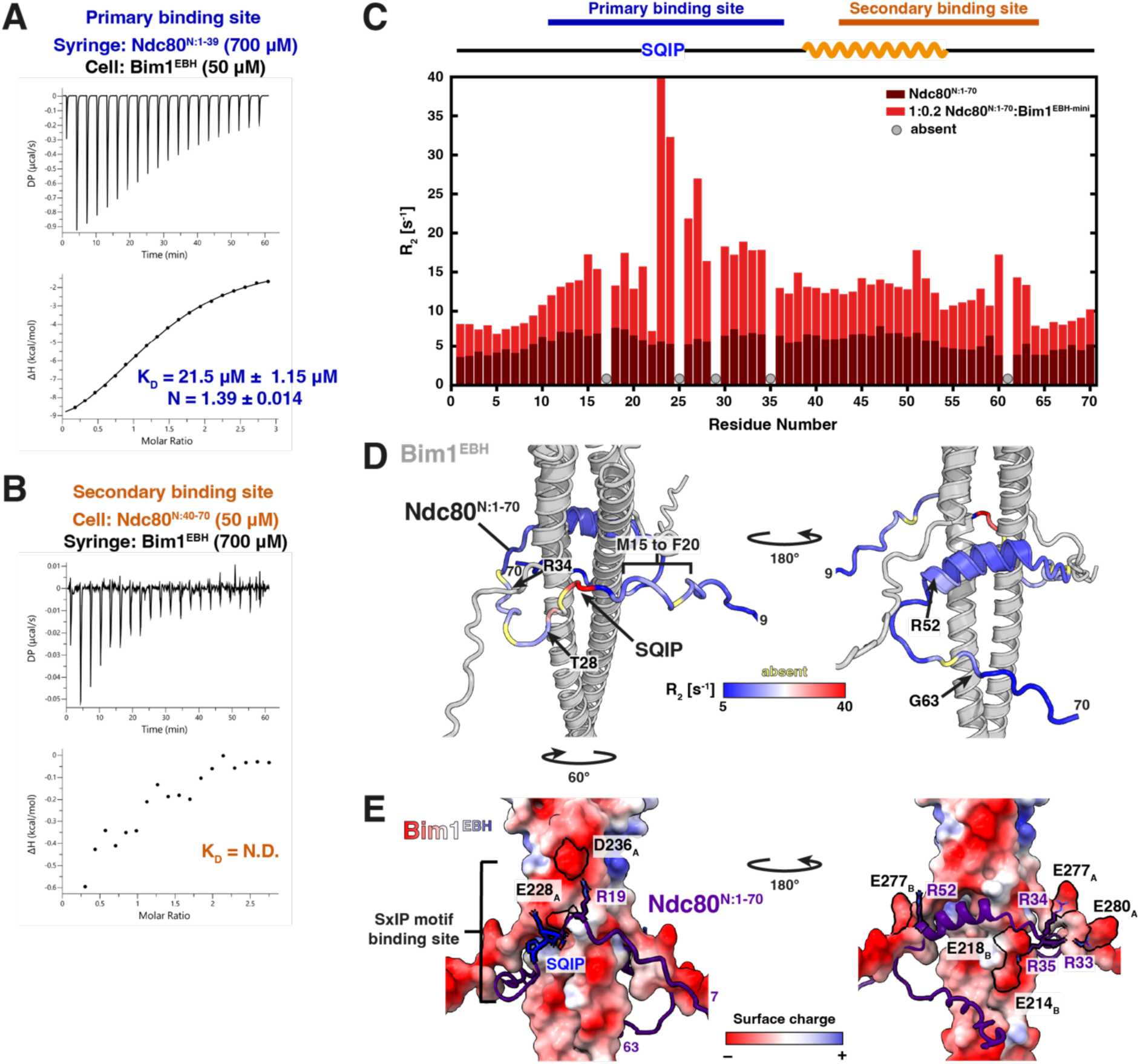
– Insights into residues contributing to the primary and secondary Bim1-binding sites of Ndc80^N^. **(A)** ITC of the primary binding site, Ndc80^N:1-39^, and Bim1^EBH^. This gave a KD of 21.5 µM and N of 1.4. **(B)** ITC of the secondary binding site, Ndc80^N:40-70^, and Bim1^EBH^. The heats upon injection indicated very weak binding that could not be fitted. **(C)** R2 (1/T2) relaxation rates of ^15^N-labelled Ndc80^N:1-70^ alone and in sub-stoichiometric complex with Bim1^EBH-mini^ (1:0.2 ratio). R2 rates are elevated in the complex due to slower tumbling and anisotropy. Residues involved and surrounding the SQIP motif (Gln23-Ala27) experience additional relaxation caused by chemical exchange (no data for Pro25). **(D)** R2 rates from (C) plotted onto the Ndc80^N:1-70^-Bim1^EBH^ AlphaFold2 prediction. Ndc80^N:1-70^ is coloured according to R2 value. The SQIP and surrounding residues have the highest R2 rates; Met15-Phe20 and Thr28-Arg34 also have higher R2 rates (left). Low occupancy of the secondary binding site at this ratio results in a modest increase of R2 rates, observed for residues 51-52 and 60-63. Arg52 might contribute to the interaction via a salt bridge (right). **(E)** Ndc80^N:1-70^-Bim1^EBH^ AlphaFold2 prediction with Bim1^EBH^ coloured by surface charge. Charged residues likely forming salt bridges are labelled. ‘A/B’ denotes different chains of Bim1^EBH^.

### The primary Bim1-binding site of Ndc80^N^ forms a canonical SxIP:EBH domain interaction

To gain further mechanistic insight into which residues within the primary binding site, in addition to the SQIP motif, contribute to Bim1–Ndc80c interactions, we performed backbone amide ^15^N T_2_ experiments (analysed as R_2_ rates). T_2_ experiments measure the dynamic properties of a given residue but are also affected by the contribution of a bound state due to the timescale of exchange between bound and unbound states typical of protein-protein interactions, thereby providing quantitative information on residues at protein interfaces (Moraes & Valente, 2023; Walinda et al., 2018). To optimise complex occupancy whilst avoiding the loss of signals observed at higher ratios, ^15^N T_2_ experiments were acquired at a 1:0.2 ratio of ^15^N-labelled Ndc80^N:1-70^ to unlabelled Bim1^EBH-mini^. Binding of Bim1^EBH-mini^ increased the overall R_2_ rates of Ndc80^N:1-70^, as expected from the slower rotational tumbling of the larger, more anisotropic complex. In addition, elevated R_2_ rates may reflect reduced local flexibility of Ndc80^N:1-70^ upon binding (Figure 5C). Residues 10-63 showed the largest increases in R_2_ rates, consistent with our titration experiments indicating that this region constitutes the Bim1-binding interface within Ndc80^N:1-70^.

At the primary binding site, residues 23-27 (the QIP residues of the SQIP motif and the adjacent TA residues) showed the most significant increase in R_2_ rates upon Bim1^EBH-mini^ binding, indicating that these residues are directly involved in the binding interface (Figure 5C, D). Because prolines lack backbone amide protons, the R_2_ rate of Pro25 could not be measured, however, based on the elevated R_2_ rates of neighbouring residues and the established role of SxIP motifs in EBH-domain interactions (Buey et al., 2012), it is likely that Pro25 contributes to binding. The low R_2_ rate of Ser22 of the SQIP motif is most likely an artefact due to partial resonance overlap with Ser6 outside of the Bim1-binding region.

Residues 15-20 (MDPHRF) and 28-34 (TSSQLRR) of the primary binding site also exhibited modest increases in R_2_ rates upon Bim1^EBH-mini^ binding, suggesting that these regions contribute to the Bim1^EBH-mini^:Ndc80^N:1-70^ interaction (Figure 5C, D). A role for residues 30-34 in Bim1^EBH^ binding is consistent with the higher affinity of Ndc80^N:1-39^ for Bim1^EBH^ compared with Ndc80^N:15-29^ (Figure 2D and Figure 5A). In the EB1:MACF1 complex, basic residues flanking the SxIP motif of MACF1 form salt bridges with negatively charged residues on EB1 (Honnappa et al., 2009). Similarly, our AlphaFold2 model predicts that Ndc80 residues Arg19, Arg33, and Arg34 form salt bridges with acidic residues surrounding the Bim1^EBH^ SxIP motif-binding site, analogous to the interactions observed in the EB1:MACF1 complex (Figure 5E). Overall, the structural features of the Bim1^EBH^:Ndc80 primary binding interface are consistent with crystal structures of other EB-family proteins bound to their partners (Honnappa et al., 2009; Matsuo et al., 2016), revealing a conserved mechanism of EBH-domain recognition involving the Ndc80 SQIP motif and its flanking residues.

Finally, consistent with both our AlphaFold2 model and NMR data, ITC analysis showed that deletion of the SQIP motif reduced the affinity of Ndc80^N:1-70^ for Bim1^EBH^ ten-fold, with Ndc80^N:1-70ΔSQIP^ binding Bim1^EBH^ with a K_D_ of 41 µM (Figure S2E, G). Together, these results demonstrate that Ndc80^N^ binds directly to the SxIP-binding site of Bim1^EBH^ through its SQIP motif.

### Structural characterisation of the secondary Bim1-binding site of Ndc80^N^

Because of the low Ndc80^N:1-70^ occupancy at the secondary binding site at low Bim1^EBH-mini^ concentrations (Figure 4C), ^15^N T_2_ experiments provided limited insight into which residues within this region of Ndc80 specifically interact with Bim1^EBH-mini^ (Figure 5C). However, elevated R_2_ rates for residues up to Gly63 and the distinct drop in R_2_ relaxation rates for residues following Gly64 indicate that residues C-terminal of Gly63 are not involved in the interaction. Furthermore, as with the SQIP motif engaging the primary binding site, we assumed that the peaks most strongly attenuated upon titration of Bim1^EBH-mini^ correspond to residues directly involved in binding (Figure 4C). This analysis identified residues 51-58 as contributors to the interaction (Figure 2A). Among these, residues 51-53 are conserved, consistent with the modestly elevated R_2_ rates observed for residues 51-52 relative to neighbouring residues. Arg52 may form a salt bridge with Bim1^EBH^ within the secondary binding interface (Figure 5D, E).

### Ipl1 phosphorylation of Ndc80 weakens the Bim1–Ndc80 interaction

Phosphorylation of Ndc80^N^ by the error correction protein kinase Ipl1 weakens kinetochore-microtubule attachments, likely through charge repulsion with negatively charged tubulin tails (Cheeseman et al., 2006; Guimaraes et al., 2008; Wei et al., 2007; Welburn et al., 2010; Wimbish & DeLuca, 2020). A recent cryo-EM study further showed that phosphorylation of human Ndc80^N^ disrupts the longitudinal oligomerisation of Ndc80 complexes on microtubules (Niu et al., 2026). The *S. cerevisiae* Ndc80^N^ incorporates seven putative Ipl1 sites (Akiyoshi et al., 2009), three of which (Thr21, Ser37 and Thr54) are within the Bim1-binding region (Figure 2A). To determine whether Ipl1 phosphorylation of Ndc80^N^ regulates the Bim1–Ndc80c interaction, we performed *in vitro* phosphorylation and monitored the resultant affinity and structural changes using ITC and NMR spectroscopy, respectively.

To assess how Ipl1 phosphorylation of Ndc80^N^ affects its affinity for Bim1, we performed ITC experiments using both phosphorylated Ndc80^N:1-70^ (pNdc80^N:1-70^) and a phosphomimetic Ndc80^N:1-70^ mutant. Phosphorylation of Ndc80^N:1-70^ caused a modest three-fold reduction in affinity for Bim1^EBH^ (K_D_ of 11.8 µM) (Figure 6A and Table S1). This is reminiscent of the phosphorylation-dependent reduction in affinity for the human EB1-APC interaction, in which the phosphorylation sites are located at distances from the SxIP motif comparable to those in Ndc80^N^ (Honnappa et al., 2005). The double phosphomimetic mutant Ndc80^N:1-70(S37E, T54E)^ bound Bim1^EBH^ with a similar affinity as pNdc80^N:1-70^ (K_D_ of 11.6 µM) (Figure 6B). Intact mass spectrometry showed that the pNdc80^N:1-70^ sample used for the ITC experiment was predominantly singly phosphorylated (70%), with only 10% doubly phosphorylated (Figure S5E). Based on the relative rates of Ser37 and Thr54 phosphorylation determined from the NMR-time course discussed below, we infer that more than 80% of pNdc80^N:1-70^ contained phosphoSer37. It is possible that fully doubly phosphorylated pNdc80^N:1-70^ would have a further reduced affinity for Bim1^EBH^. Consistent with this, single phosphomimetic mutants bound Bim1^EBH^ more tightly than the double phosphomimetic mutant, with K_D_ values of 7.4 µM and 7.7 µM measured for Ndc80^N:1-70(S37E)^ and Ndc80^N:1-70(T54E)^, respectively (Figure S5C, D and Table S1). These results indicate that, despite their different distances from the SQIP motif, phosphorylation at Ser37 and Thr54 contributes similarly to reducing the affinity of Ndc80^N^ for Bim1^EBH^.

**Figure 6.**
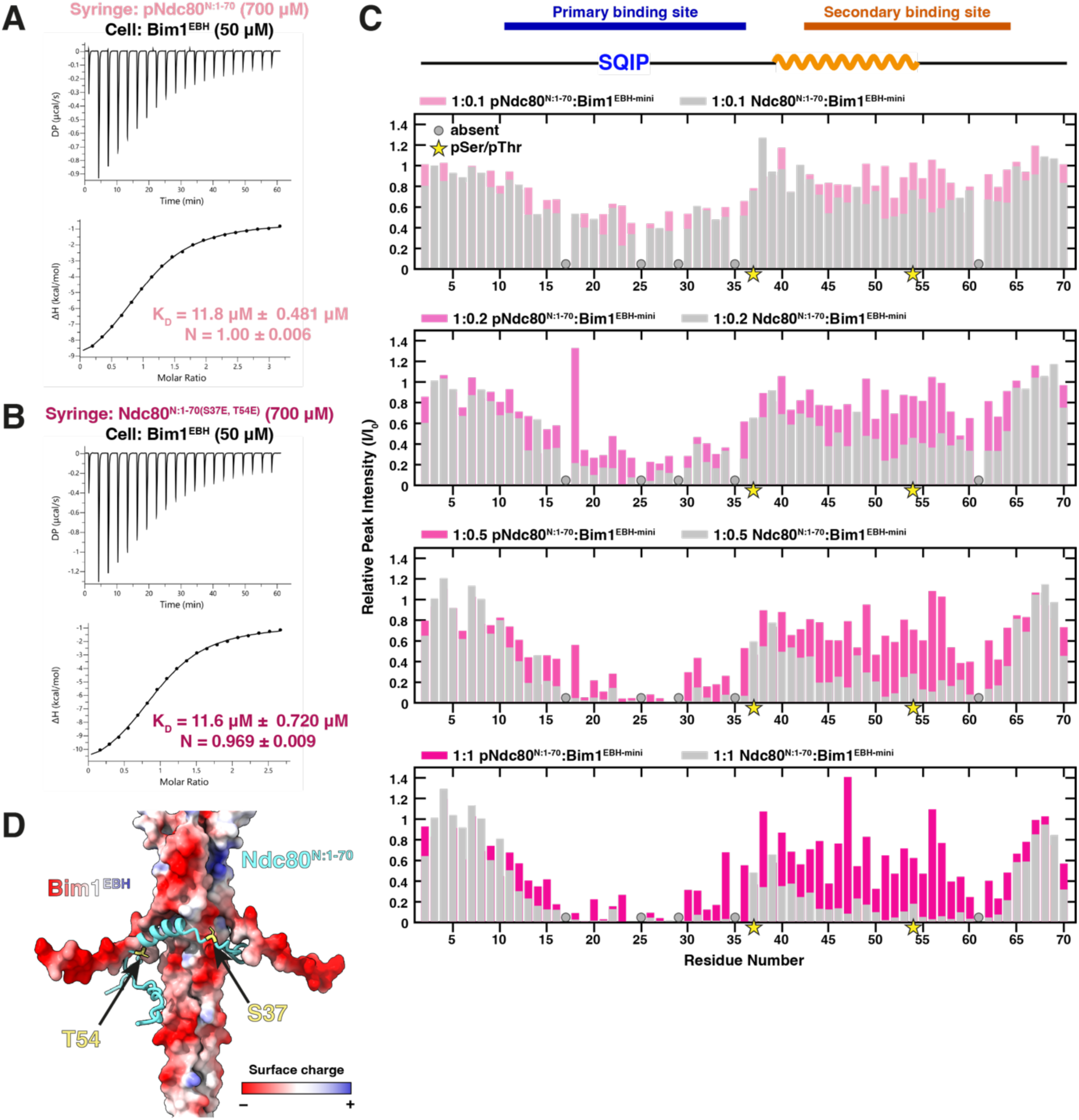
– Ipl1 phosphorylation of Ndc80^N^ weakens the Bim1-Ndc80c interaction. **(A)** ITC of pNdc80^N:1-70^ with Bim1^EBH^. This gave a KD of 11.8 µM and N of 1. **(B)** ITC of Ndc80^N:1-70(S37E, T54E^) with Bim1^EBH^. This gave a KD of 11.6 µM and N of 1. **(C)** Relative peak intensities (I/I0) of peaks from the 2D ^1^H, ^15^N HSQC spectra of ^15^N-labelled pNdc80^N:1-70^ titrated with the indicated ratios of Bim1^EBH-mini^ overlayed with the relative peak intensities for the same titration ratio of Ndc80^N:1-70^ with Bim1^EBH-mini^ (from Figure 4C). There is generally less peak attenuation in the presence of Bim1^EBH-mini^ for the Bim1-binding regions of pNdc80^N:1-70^ than Ndc80^N:1-70^. The secondary binding site around Ndc80^N-helix^ is more significantly affected by phosphorylation. Intensities are normalised relative to Ser3. **(D)** AlphaFold2 prediction of the Bim1^EBH^-Ndc80^N:1-70^ interaction with Bim1^EBH^ coloured by surface charge, highlighting Ser37 and Thr54. Ser37 and Thr54 are predicted with low confidence (Figure 3), however our NMR data agree with the prediction.

### Ipl1 phosphorylation changes the structure and dynamics of Ndc80^N^

To monitor changes in Ndc80^N:1-70^ conformation caused by Ipl1 phosphorylation, we collected 2D ^1^H,^15^N HSQC spectra of ^13^C,^15^N-labelled Ndc80^N:1-70^ at 10 min intervals. To slow the phosphorylation reaction sufficiently for time-resolved analysis, we used a 1:5000 ratio of the Ipl1/Aurora B kinase complex (Ipl1-Sli15) to Ndc80^N:1-70^ (Figure 7A and Figure S5A). Phosphorylation causes characteristic ^1^H downfield shifts of Ser/Thr residues from 7.8-8.2 ppm (Ser/Thr region) to 8.5-8.8 ppm (pSer/pThr region) (Bienkiewicz & Lumb, 1999). A signal corresponding to Ser37 phosphorylation was first detected approximately 1.5 h after kinase addition, whereas Thr54 phosphorylation was first observed after ∼5 h (Figure 7A). The faster phosphorylation of Ser37 is consistent with the preference of Ipl1/Aurora B for motifs containing basic residues at the -2 and -3 positions (Figure 2A) (Meraldi et al., 2004), and may reflect the relative order of phosphorylation *in vivo*. In multiple repeats, we did not observe Thr21 phosphorylation, consistent with phosphoproteomic data showing phosphorylation of Ser37 and Thr54, but not Thr21 *in vivo* (Akiyoshi et al., 2009). These data indicate that Thr21 is not efficiently phosphorylated by Ipl1 despite matching the consensus sequence, possibly because Pro17 at the -4 position induces structural features that interfere with kinase recognition of Ndc80.

**Figure 7.**
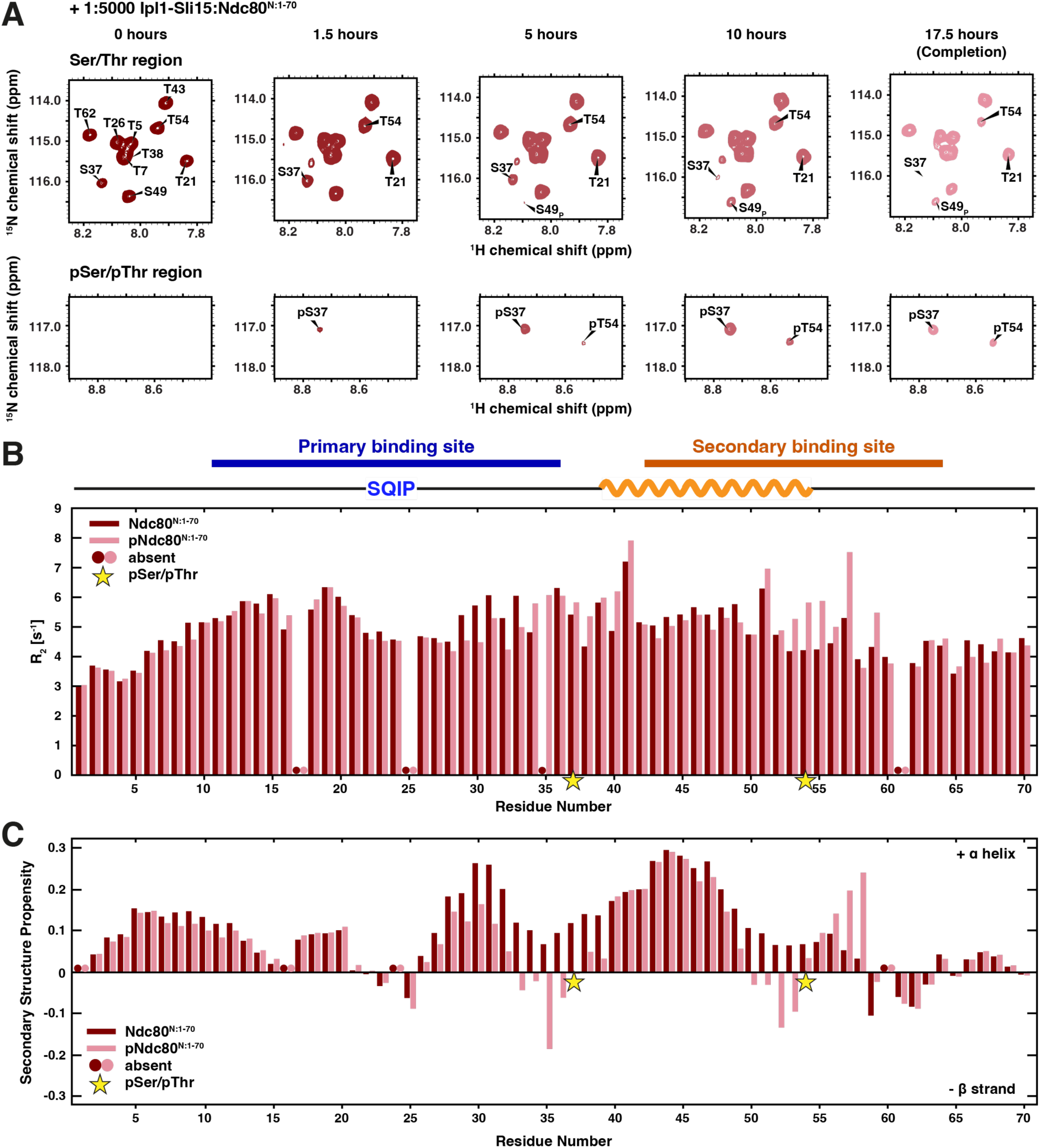
– Ipl1 phosphorylation of Ser37 and Thr54 of Ndc80^N^ changes its secondary structure propensity. **(A)** Expanded Ser/Thr (top) and pSer/pThr (bottom) regions of 2D ^1^H, ^15^N HSQC spectra of ^13^C,^15^N-labelled Ndc80^N:1-70^ with a ratio of 1:5000 Ipl1-Sli15:Ndc80^N:1-70^ at the specified time points after kinase addition. Spectra were collected roughly every 10 min to track changes due to phosphorylation. pSer37 appears before pThr54, Thr21 is not phosphorylated. Ser49 experiences a shift due to change in chemical environment (S49P). **(B)** R2 rates of ^15^N-labelled Ndc80^N:1-70^ and pNdc80^N:1-70^ (two bars per residue). **(C)** Secondary structure propensities of ^13^C,^15^N-labelled Ndc80^N:1-70^ and pNdc80^N:1-70^ (two bars per residue), based on differences between experimental and random coil chemical shifts accounting for phosphorylated residues. Residues preceding prolines (‘absent’) are not considered.

Calculation of weighted chemical shift perturbations (CSPs) between Ndc80^N:1-70^ and Ser37/Thr54 phosphorylated Ndc80^N:1-70^ (pNdc80^N:1-70^) showed that residues 34-36 and 49-56 undergo significant perturbations, whereas all other residues remained at the same chemical shift position (Figure S5A, B). These phosphorylation-dependent CSPs could result either from the direct addition of a nearby phosphate group, or from phosphorylation-induced changes in the dynamics and secondary structure propensity of Ndc80^N:1-70^. To distinguish between these possibilities, we performed ^15^N T_2_ experiments and calculated secondary chemical shifts for pNdc80^N:1-70^ using theoretical random coil chemical shifts that account for phosphorylation (Hendus-Altenburger et al., 2019). Phosphorylation of Ndc80^N:1-70^ reduced R_2_ relaxation rates of residues 29-34 within the primary binding site, indicating that Ser37 phosphorylation increases the conformational flexibility of this region in solution (Figure 7B). This change correlated with a reduction in α-helical propensity (Figure 7C). Within the secondary binding site, phosphorylation at Thr54 destabilised α-helical propensity at the C-terminus of Ndc80^N-helix^ (residues 50-53) (Figure 7C), whereas residues 53-58 showed increased R_2_ rates, indicating that pThr54 enhances their rigidity (Figure 7B). Consistent with this observation, residues 55-58 displayed increased α-helical propensity upon phosphorylation, forming a new region that samples α-helical structure (Figure 7C). These phosphorylation-dependent changes in the dynamics of Ndc80^N^ may also contribute to the reduced microtubule-binding activity of phosphorylated Ndc80c. Interestingly, the behaviour of the SQIP motif and its immediately surrounding residues remained largely unchanged following Ipl1 phosphorylation.

### Ipl1 phosphorylation of Ndc80^N^ disrupts interaction of its primary and secondary-binding sites with Bim1

To determine how Ipl1 phosphorylation specifically affects interactions of the primary and secondary binding sites of Ndc80^N:1-70^ with Bim1^EBH^, we titrated Bim1^EBH-mini^ into ^15^N-labelled pNdc80^N:1–70^ (Figure 6C). Compared with unphosphorylated Ndc80^N:1-70^, residues of both binding sites exhibited reduced peak attenuation in the presence of Bim1^EBH-mini^, consistent with weaker binding. Although attenuation of the SQIP motif and its surrounding residues was modestly reduced, the most pronounced effects were observed at the C-terminal region of the primary binding site and throughout the secondary binding site. These differences became more apparent at higher concentrations of Bim1^EBH-mini^.

Within the primary binding site, Ser37 phosphorylation weakened the interaction of residues 31-36 with Bim1^EBH-mini^, matching the phosphorylation-dependent changes in R_2_ relaxation rates and secondary structure propensity (Figure 7B, C). Previous studies on the EBI:APC complex suggested that nearby phosphorylation weakens the contribution of salt bridges to SxIP-mediated interactions (Honnappa et al., 2009). Therefore, phosphorylation of Ser37 would be expected to reduce the electrostatic contributions of Arg33 and Arg34 to the Bim1–Ndc80c interaction (Figure 5D, E). In addition, the AlphaFold2 model predicts that Ser37 lies adjacent to a negatively charged surface patch on Bim1^EBH^ (Figure 6D), suggesting that phosphorylation at this site could further weaken binding through electrostatic repulsion. Phosphorylation of Thr54 reduced the affinity of the entire secondary binding site of Ndc80^N:1-70^ for Bim1^EBH-mini^ (Figure 6C). Because there is no exchange between the phosphorylated and non-phosphorylated species, all peaks used for analysis were from the phosphorylated species. Phosphorylation of Thr54 at the C-terminus of Ndc80^N-helix^ weakened the contribution of this region to Bim1 binding (Figure 5C), likely through destabilisation of Ndc80^N-helix^ (Figure 7C), disruption of Arg52-dependent electrostatic interactions, and electrostatic repulsion with negatively charged regions of Bim1^EBH^ (Figure 6C,D). The C-terminal region of the secondary binding site also showed reduced binding to Bim1^EBH-mini^ following phosphorylation (Figure 6C), consistent with the substantial phosphorylation-dependent changes in secondary structure propensity observed for this region (Figure 7C).

In addition to Ipl1 phosphorylation, Ndc80^N^ contains Mps1 kinase phosphorylation sites (Figure 2A) (Kemmler et al., 2009). Mps1 is the master regulator of the spindle assembly checkpoint (SAC) and has been implicated in the error-correction mechanism. Phosphorylation of Ndc80^N^ by Mps1 weakens kinetochore–microtubule attachments *in vitro* (Hayward et al., 2022; Kops & Shah, 2012; Sarangapani et al., 2021). To examine how Mps1-dependent phosphorylation affects the affinity of Ndc80^N:1-70^ for Bim1, we generated an Mps1 phosphomimetic mutant, Ndc80^N:1-70(Mps1)^, in which all five Mps1 phosphorylation sites within the Bim1-binding region were replaced with glutamate (T21E, S22E, S37E, T38E, T43E). ITC analysis showed that this mutant binds Bim1^EBH^ with a greatly reduced affinity, exhibiting a K_D_ of 142 µM, an approximately 35-fold decrease relative to Ndc80^N:1-70^ (Figure S5F and Table S1). These results suggest that Mps1-dependent phosphorylation could regulate the Bim1–Ndc80c interaction during error correction.

### Bim1 augments Ndc80c-microtubule attachment strengths *in vitro*

To explore how the Bim1–Ndc80c interaction affects the strength of Ndc80c-microtubule attachments, we used an *in vitro* optical trap rupture force assay which measures the force required to detach reconstituted kinetochore complexes from dynamic microtubules (Figure S6A-C). We determined that the median rupture force for the Ndc80c-microtubule attachment was 5.2 pN, consistent with previous results (Muir et al., 2023; Tien et al., 2010; Umbreit et al., 2014). Addition of Bim1 increased this rupture force to 8.7 pN (Figure 8A), indicating that the Bim1:Ndc80c complex forms in the context of microtubules and enhances the load-bearing capacity of Ndc80c attachments.

**Figure 8.**
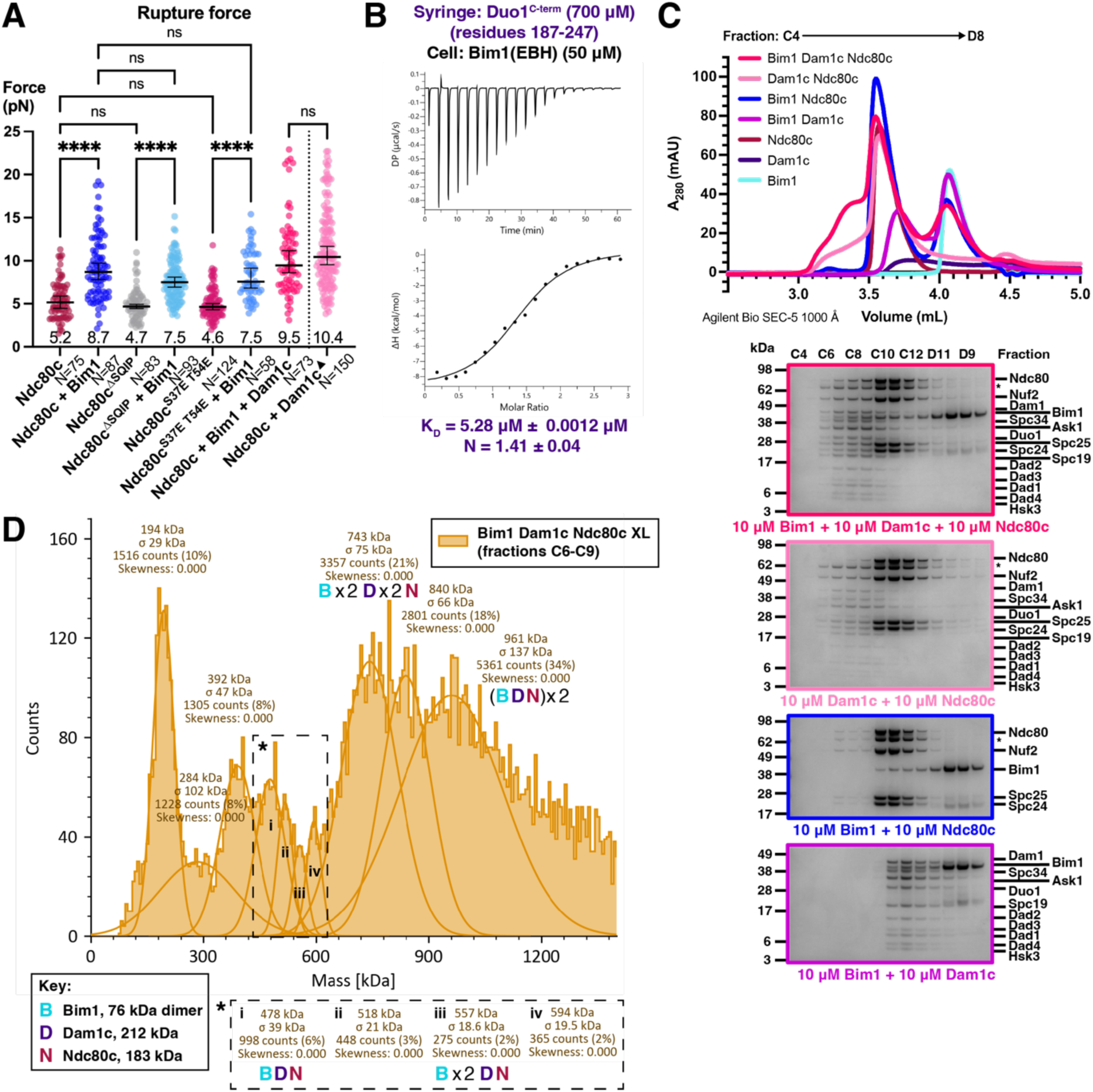
– Bim1-Ndc80c interaction in the context of outer kinetochore-microtubule attachments. **(A)** Optical trap rupture force measurements of Ndc80c-microtubule attachments without and with Bim1. Ndc80c, Ndc80c^ΔSQIP^ or Ndc80c^S37E^ ^T54E^ were immobilised on streptavidin-coated beads, Bim1 and Dam1 were added to the flow cell as indicated. Each dot represents a single rupture event measurement. Black bar shows the median rupture force values (number stated below) with 95% confidence intervals over N measurements. A Kruksal-Wallis test was performed to determine statistical significance of the difference in medians. **** = P value < 0.0001; ns, not significant. ^▴^ Ndc80c + Dam1c dataset from (Muir et al., 2023). **(B)** ITC of Duo1^C-term^ with Bim1^EBH^. This gave a KD of 5.28 µM and N of 1.4. **(C)** Chromatogram (top) and corresponding SDS-PAGE analysis (bottom) of eluate fractions of all combinations of Bim1, Ndc80c and Dam1c injected onto an Agilent Bio SEC-5 1000 Å column individually and together at equimolar. Bim1, Ndc80c and Dam1c alone gels are shown in Figure S8D. * = Ndc80^ΔN^ where some degradation of Ndc80^N^ occurred during purification. **(D)** Stacked histograms of mass photometry measurements taken of a crosslinked Bim1-Dam1c-Ndc80c sample after SEC as in part (C) (fractions C6-C9). The molecular masses, error, and population of each mass class are indicated. Possible stoichiometries of observed complexes likely to comprise all three of Bim1, Dam1c and Ndc80c are indicated below the molecular mass for the class.

We next performed rupture force assays with Ndc80^N^ mutants designed to disrupt the Bim1–Ndc80c interaction while minimising the effects of Ndc80^N^-microtubule binding. Ndc80c^ΔSQIP^ and Ndc80c^S37E, T54E^ displayed median rupture forces of 4.7 pN and 4.6 pN, respectively, and rupture force distributions similar to wild-type Ndc80c (Figure 8A). In the presence of Bim1, the median rupture forces of both Ndc80c^ΔSQIP^ and Ndc80c^S37E, T54E^ increased to 7.5 pN. Relative to wild-type Ndc80c with Bim1, this is a modest reduction in rupture force, and although fewer high-force rupture events were observed, these differences were not statistically significant (Figure 8A). As shown above (Figure S2G and Table S1), although the SQIP motif is a major determinant of the Bim1–Ndc80c interaction, its deletion still permits binding to Bim1. Ndc80c retains microtubule-binding activity upon Ipl1 phosphorylation, with error correction involving phosphorylation of multiple outer kinetochore components including Ndc80 and Dam1 (Akiyoshi et al., 2009; Cheeseman et al., 2002; Doodhi et al., 2021; Muir et al., 2023). Dam1 is likely to be the more important Ipl1 target in mediating error correction. We previously showed that phosphomimetic mutations of Ipl1 target sites in Dam1 abolishes the ability of Dam1c to enhance Ndc80c–microtubule attachment strength (Muir et al., 2023).

Bim1 and Ndc80c independently interact with Dam1c (Dudziak et al., 2021; Kim et al., 2017; Muir et al., 2023). In our rupture force assay, simultaneous addition of Bim1 and Dam1c produced a median Ndc80c-microtubule rupture force of 9.5 pN (Figure 8A), similar to the 10.4 pN observed previously with Dam1c alone (Muir et al., 2023). These results indicate that the Ndc80c-Dam1c interaction provides the dominant contribution to kinetochore-microtubule attachment strength, and that Bim1 and Dam1c do not cooperate to further strengthen Ndc80c-microtubule attachments. The basis for this lack of enhancement is unclear. One possibility is that the maximum rupture force measured in these optical trap assays is limited by the load-bearing capacity of the Ndc80c:Dam1c assembly itself, rather than by the strength of the Ndc80c-microtubule interaction. Using ITC we found that the C-terminus of the Dam1c subunit Duo1 (Duo1^C-term^), which contains an SxIP motif, bound Bim1^EBH^ with a K_D_ of 5.3 µM and a stoichiometry of 1.4 (Figure 8B and Table S1). This affinity is higher than either the primary or secondary Bim1-binding sites of Ndc80c individually. In addition, the Dam1c subunits Spc19 and Spc34 may also contribute to the Bim1-Dam1c interaction (Dudziak et al., 2021). Thus, it is likely that Dam1c competes with Ndc80c for Bim1 binding.

Because microtubule binding by Ndc80c and Dam1c is compatible with Bim1 association, we investigated the interplay between Bim1, Dam1c, and Ndc80c using analytical SEC. When mixed at equimolar concentrations of 10 µM, Bim1, Dam1c, and Ndc80c, Bim1 co-eluted together with Dam1c and Ndc80c, indicating formation of a Bim1:Dam1c:Ndc80c ternary complex (Figure 8C). To further characterise this assembly, we performed mass photometry on crosslinked Bim1:Dam1c:Ndc80c complexes following SEC purification. The resulting heterogeneous molecular mass distribution indicated the presence of multiple distinct species, likely corresponding to different stoichiometries of the Bim1:Dam1c:Ndc80c complex as well as pairwise subcomplexes (Figure 8D). These observations are consistent with the low-affinity, dynamic interactions characteristic of EB-family proteins, which are thought to facilitate continual remodelling of protein networks at microtubule plus ends (Akhmanova & Steinmetz, 2008, 2015). It is therefore possible that multiple combinations of Bim1, Dam1c, and Ndc80c interactions coexist during the establishment of end-on kinetochore–microtubule attachments.

### *In vitro* reconstitution of Bim1–Ndc80c binding to dynamic microtubules

Bim1 autonomously tracks microtubule plus-ends (Howes et al., 2018; Lampert et al., 2010; Zimniak et al., 2009), whereas Ndc80c lacks this activity (Lampert et al., 2010). Because the Bim1-Dam1c interaction recruits Dam1c to microtubule plus-ends (Dudziak et al., 2021), we tested whether Bim1 could similarly recruit Ndc80c. We therefore used a TIRF microscopy-based assay to reconstitute Ndc80c binding to dynamic microtubules in either the presence or absence of Bim1 (Figure S7). 150 mM KCl was used in the imaging buffer to reduce Ndc80c binding to the microtubule lattice (Lampert et al., 2010). Without Bim1, we observed no binding of Ndc80c-SNAP-Alexa Fluor 647 (Ndc80c-647) to dynamic microtubules (n = 122) (Figure S7). Upon addition of unlabelled Bim1, we observed binding of Ndc80c-647 dynamic microtubules (n = 167 of 269 microtubules), consistent with our optical trap assays (Figure S7 and Figure 8A). We additionally observed a limited population of Ndc80c-647 colocalised with microtubule plus-ends (n = 4 of 269 microtubules) (Figure S7), suggesting that Bim1 could localise Ndc80c to microtubule plus-ends. In support of this, the *Schizosaccharomyces pombe* Bim1/EB1 homologue Mal3 confers microtubule plus-end tracking activity to Ndc80c *in vitro*, however, the molecular basis of the Mal3-Ndc80 interaction is not known (Matsuo et al., 2017). The rare, short-lived nature of Bim1-mediated Ndc80c tip tracking observed here is consistent with Bim1 being inefficient at tip tracking on vertebrate microtubules (Howes et al., 2018; Molodtsov et al., 2016). Bim1 robustly tracks plus-ends of yeast microtubules *in vitro* (Howes et al., 2018), however, due to technical limitations we were unable to perform assays with yeast microtubules.

### Disruption of the Bim1-binding interface does not compromise chromosome segregation

Unlike in higher eukaryotes, the N-terminal region of Ndc80 is not essential in budding yeast (Akiyoshi et al., 2009; Guimaraes et al., 2008; Kemmler et al., 2009; Miller et al., 2008). Nevertheless, disruption of multiple Ipl1 or Mps1 phosphorylation sites within Ndc80^N^ confers benomyl sensitivity, indicating an important role for this region in kinetochore function (Akiyoshi et al., 2009; Kemmler et al., 2009). To investigate the regulation of the Bim1–Ndc80c interaction *in vivo*, we generated yeast strains for ectopic expression of Ndc80^N^ mutants in an Ndc80-AID background (Muir et al., 2023) (Figure S8), in which endogenous Ndc80 can be depleted using an auxin-inducible degron system (Tanaka et al., 2015). In the absence of wild-type Ndc80, cells expressing either Ndc80^ι1N^ or Ndc80^S37A,T54A^ were resistant to benomyl (Figure S8), consistent with previous observations (Akiyoshi et al., 2009; Demirel et al., 2012). Similarly, cells expressing Ndc80 with either a disrupted SQIP motif (Ndc80^ΔSQIP^) or the phosphomimetic mutations S37E and T54E exhibited normal growth on benomyl, indicating that perturbation of the Ndc80^N^-Bim1 interface by disruption of the SQIP motif or Ser37/Thr54 phosphorylation does not compromise chromosome segregation. These *in vivo* results are consistent with our rupture-force measurements, which showed that the ability of Bim1 to strengthen Ndc80-microtubule attachments is unaffected by disruption of the SQIP motif or by the S37E,T54E phosphomimetic mutations. Furthermore, cells expressing phosphonull or phosphomimetic mutations of all Mps1 phosphorylation sites in the Bim1-binding region of Ndc80^N^ (Thr21, Ser22, Ser37, Thr38, Thr43) exhibited normal growth on benomyl, consistent with previous observations that mutation of Mps1 sites within both Ndc80^N^ and the Ndc80 CH domain is required to disrupt cell growth on benomyl (Kemmler et al., 2009).

## Discussion

In this study we identified and characterised a direct interaction between the +TIP protein Bim1 and the outer kinetochore Ndc80 complex in *S. cerevisiae*. We showed that this interaction is mediated primarily through recognition of a conserved SQIP motif within the intrinsically disordered N-terminus of Ndc80 by the Bim1 EBH domain. The Ndc80 SQIP motif alone is insufficient to confer full interaction affinity, and a secondary binding site C-terminal to the SQIP motif increases interaction strength nearly 50-fold. We also present the first NMR analysis of Ndc80^N^, revealing that it contains segments that adopt α-helical structure, the largest region being within the secondary Bim1-binding site. Phosphorylation of Ser37 and Thr54 by Ipl1 substantially alters the dynamic behaviour of Ndc80^N^ and weakens its interaction with Bim1. Ndc80c interaction with Bim1 increases the strength of Ndc80-microtubule attachments but does not further increase the strength of Ndc80c-Dam1c-microtubule attachments, even though Bim1, Dam1c and Ndc80c form a ternary complex. Electron microscopy and CL-MS support a model in which a Bim1 dimer bridges two Dam1c protomers via interactions between the Duo1 subunit-SxIP motif and the Bim1 EBH domain (Dudziak et al., 2021). Although this interaction is displaced by Ndc80c (Dudziak et al., 2021), our findings reveal that Bim1 still interacts with Ndc80c when Dam1c is present. This observation suggested the possibility that Bim1 could enhance Ndc80c:Dam1c–microtubule attachment strength. However, our rupture force experiments showing that Bim1 and Dam1c do not cooperatively strengthen Ndc80c–microtubule attachments, do not support this idea.

We therefore propose a model in which the major function of the Bim1–Ndc80c interaction is to facilitate localisation of Ndc80c to microtubule plus-ends during the establishment of end-on kinetochore-microtubule attachments, either by recruiting Ndc80c to plus-ends and/or by stabilising Ndc80c at these sites (Figure 9). Ndc80c does not autonomously track microtubule plus-ends, and the number of Ndc80 complexes at kinetochores increases from G1 to anaphase (Dhatchinamoorthy et al., 2017; Lampert et al., 2010). Bim1 may therefore contribute to the initial recruitment of Ndc80c to plus-ends and maintain its attachment prior to formation of mature, Dam1c-mediated end-on kinetochore-microtubule attachments. Bim1 has a similar role in localising Dam1c, and additionally promotes Dam1c oligomerisation (Dudziak et al., 2021). Supporting this proposed role of *S. cerevisiae* Bim1 in recruiting Ndc80c to plus-ends, we observed some Bim1-mediated localisation of Ndc80c to dynamic microtubule plus-ends by TIRF microscopy. Additionally, the *S. pombe* Bim1/EB1 homologue Mal3 confers microtubule plus-end tracking activity on Ndc80c *in vitro* (Matsuo et al., 2017). However, the molecular basis of the Mal3-Ndc80 interaction was not defined (Matsuo et al., 2017), and *S. pombe* Ndc80 lacks canonical EBH-domain-binding motifs. On formation of incorrect attachments, Ipl1 phosphorylation activity weakens the Bim1–Ndc80c interaction (Figure 5A, B), simultaneously suppressing Ndc80c-microtubule binding, the Ndc80c-Dam1c interaction, and Dam1c ring oligomerisation (Figure 9) (Akiyoshi et al., 2009; Cheeseman et al., 2002; Doodhi et al., 2021; Muir et al., 2023; Wei et al., 2007). Also consistent with a role of Bim1 in establishing, rather than maintaining end-on kinetochore-microtubule attachments, its phosphorylation by Ipl1 did not affect Bim1’s capacity to strengthen Ndc80-microtubule attachment strength.

**Figure 9.**
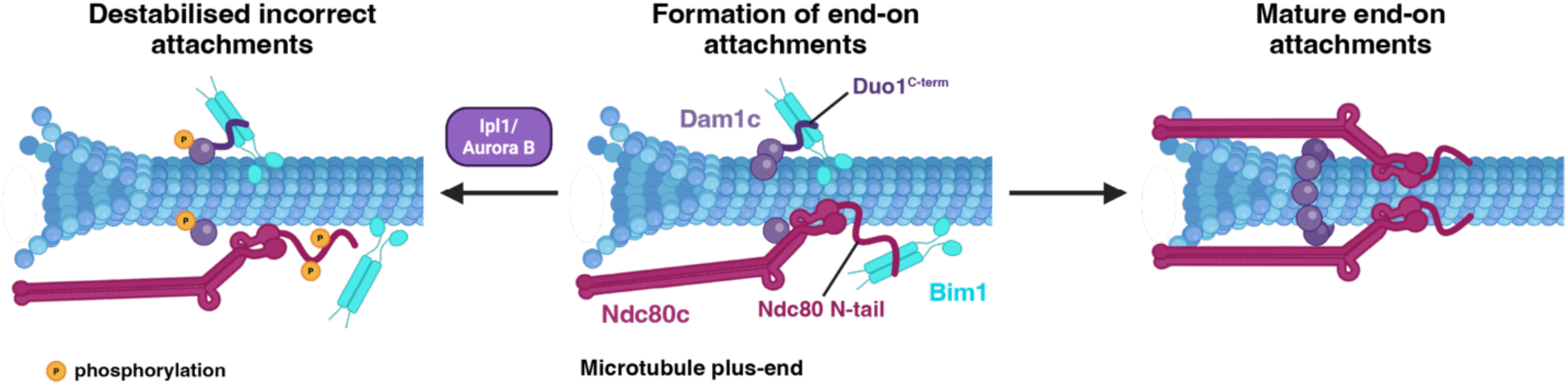
– Model for the role of the Bim1-Ndc80c in kinetochore-microtubule attachments. Bim1 is localised at microtubule plus-ends and binds Ndc80c (via the Ndc80 N-tail) and Dam1c (via the Duo1 C-terminus), localising them during the formation of end-on attachments. Bim1 also promotes Dam1c oligomerisation, and initial Ndc80c-Dam1c interactions form (middle). Incorrect, low-tension kinetochore-microtubule attachments are phosphorylated by Ipl1 (Aurora B), which weakens the Bim1-Ndc80c interaction and promotes dissociation of Dam1c oligomers (left). Stable Ndc80c-Dam1c interactions occur with formation of the Dam1c ring, giving mature end-on kinetochore-microtubule attachments (right). Created with BioRender.com.

The low micromolar-affinity of the Bim1–Ndc80c interaction *in vitro* is comparable with other +TIP-mediated interactions, whose relatively low affinities are thought to facilitate rapid remodelling of the highly dynamic +TIP network at microtubule plus-ends (Akhmanova & Steinmetz, 2008, 2015). We have furthermore shown that the interaction occurs at nanomolar concentrations in the context of microtubules.

The mechanism underlying the Bim1–Ndc80c interaction described here is distinct from previously characterised EBH-mediated interactions. In this case, a secondary binding site lacking a canonical SxIP or LxxPTPh motif engages the second SxIP-binding pocket of the Bim1^EBH^ dimer. Previous studies of SxIP-mediated interactions have revealed how residues immediately flanking the SxIP motif contribute to binding (Almeida et al., 2024; Honnappa et al., 2009; Kumar et al., 2017; Matsuo et al., 2016), and have shown that many EB-family binding partners contain multiple SxIP or LxxPTPh motifs to enhance interaction affinity (Akhmanova & Steinmetz, 2008; Applewhite et al., 2010; Kumar et al., 2017; van der Vaart et al., 2011; Zimniak et al., 2009). Consistent with these earlier studies, our model suggests that positively charged residues within both the primary and secondary binding sites contribute to binding through electrostatic interaction with the negatively charged EBH domain (Honnappa et al., 2009). Using NMR spectroscopy, we obtained mechanistic insight into the specific residues that stabilise the Bim1:Ndc80c complex.

The presence of an independent secondary binding site that wraps around the EBH domain suggests a role beyond simply increasing interaction affinity. One possible function is to act as a stoichiometry determinant, ensuring only a single Ndc80c binds each Bim1 dimer. Limiting the stoichiometry of the interaction may prevent an excessive accumulation of Bim1-bound Ndc80c at microtubule plus-ends. Furthermore, the secondary binding site may function in the error correction pathway by disrupting the localisation of Ndc80c to microtubule plus-ends during the establishment of end-on kinetochore-microtubule attachments. Ipl1 preferentially weakens Bim1 binding at the Ndc80 secondary binding site. This could allow an SxIP motif from another, non-phosphorylated Ndc80c (or Dam1c) molecule to bind to Bim1, thereby facilitating the subsequent formation of a correct kinetochore-microtubule attachment.

In this study, we observed substantial changes in the secondary structure of Ndc80^N^ upon Ipl1 phosphorylation. These changes could themselves affect Ndc80^N^ contribution to Ndc80c-microtubule binding. Our data indicate that the reduced affinity of phosphorylated Ndc80^N^ for Bim1 results from a combination of secondary structure changes, weaker electrostatic interactions, and charge repulsion with negatively charged regions of the Bim1 EBH domain. These findings suggest that phosphorylation of Ndc80^N^ during error correction may disrupt kinetochore-microtubule attachments both directly through electrostatic repulsion between phosphorylated Ndc80 residues and the negatively charged tubulin tails, and indirectly by disrupting Bim1 binding (Akiyoshi et al., 2009; Cheeseman et al., 2002; Doodhi et al., 2021).

Mps1 phosphorylation of Ndc80^N^ may also regulate the Bim1–Ndc80c interaction, potentially as part of the proposed role of Mps1 in the error correction mechanism (Hayward et al., 2022; Kops & Shah, 2012; Sarangapani et al., 2021). In contrast, Mps1 phosphorylation of Dam1c outside of its Bim1-binding region promotes the Bim1-Dam1c interaction (Dudziak et al., 2021), consistent with the role of Mps1 in promoting biorientation and coupling of kinetochores to microtubule plus-ends (Maure et al., 2007; Shimogawa et al., 2006).

The human Ndc80^N^ lacks a canonical motif that would be recognised by the EB1 EBH domain, suggesting that the interaction described here is unlikely to be conserved between budding yeast and human. This observation provides further insight into the evolutionary divergence of the microtubule-coupling mechanisms between budding yeast and human outer kinetochores. Our study identifies a previously uncharacterised direct interaction between a +TIP protein and the kinetochore. The +TIP network is highly dynamic and governed by a hierarchy of numerous weak interactions (Akhmanova & Steinmetz, 2015), with hundreds of proteins localising to microtubule plus-ends during chromosome segregation, particularly in higher eukaryotes (McAinsh & Marston, 2022). Future studies unveiling the interplay between the +TIP network and the kinetochore will provide a full understanding of the mechanisms ensuring faithful chromosome segregation in cells.

## Methods

### Cloning, expression and purification of Bim1

The Bim1 coding sequence was cloned from *S. cerevisiae* S288c genomic DNA into a modified pF1 vector containing an N-terminal 6xHis-SNAP tag (3C cleavable) (Zhang et al., 2016). A baculovirus was generated from this plasmid as described (Zhang et al., 2016). Following P3 baculovirus generation, High Five insect cells were infected with 2.5% vol/vol of P3 virus and harvested when cell viability had dropped to around 80%.

All purification steps in this study were performed at 4°C or on ice using an AKTA Pure HPLC system (Cytiva) for all chromatography steps. Cells expressing Bim1 were harvested by centrifugation and lysed by sonication in lysis buffer (50 mM Tris pH 8.0, 300 mM NaCl, 5 mM benzamidine, 1 mM β-mercaptoethanol, 5% glycerol) containing 10 µL Benzonase Nuclease (Merck), 1 mM PMSF and cOmplete EDTA-free protease inhibitor tablets (Roche). Lysate was clarified by centrifugation for 30 min at 38,000 x g and loaded onto 3 x 5 mL HiTrap TALON crude columns (Cytiva) in lysis buffer, washed with lysis buffer + 10 mM imidazole, and eluted with lysis buffer + 300 mM imidazole. Eluate was diluted 1:1 with IEX buffer (20 mM Tris pH 8.0, 1 mM TCEP) to give 150 mM NaCl, incubated with 3C protease at 4°C overnight then loaded onto a RESOURCE Q column (Cytiva), washed with IEX buffer + 150 mM NaCl and eluted with a 20 CV gradient to IEX buffer + 1 M NaCl. Bim1-containing fractions were concentrated with an Amicon Ultra 50 kDa MWCO centrifugal filter (Merck Millipore) and loaded onto a HiLoad 16/600 Superdex 200 pg SEC column (Cytiva) equilibrated in SEC200 buffer (20 mM HEPES pH 7.5, 200 mM NaCl, 1 mM TCEP). Desired fractions from the SEC peak were concentrated, flash frozen in liquid nitrogen and stored at -80°C.

### Cloning, expression and purification of Ndc80c, Ndc80c mutants and Dam1c

Ndc80c and Dam1c were cloned, expressed and purified as described in (Muir et al., 2023).

Coding sequences for desired Ndc80 mutants in Ndc80c were cloned from a modified pF1 vector containing Ndc80 and Spc25 (Zhang et al., 2016). Nuf2 (or Nuf2 with a C-terminal SNAP tag for SNAP-tagged Ndc80c mutants) and Spc25 were cloned into a separate pF1 vector. Separate baculoviruses were made from these plasmids as described (Zhang et al., 2016). Following P3 baculovirus generation, High Five insect cells were co-infected with 2.5% vol/vol of each baculovirus and harvested when cell viability had dropped to around 80%. Ndc80c mutants and SNAP-tagged Ndc80c mutants were then purified as described in (Muir et al., 2023).

### Cloning, expression and purification of Ndc80^N^ constructs and Duo1^C-term^

Coding sequences from *S. cerevisiae* S288c for Ndc80^N^ constructs and Duo1^C-term^ (residues 187-247) were cloned into a modified pETM11 vector containing an N-terminal 6xHis-SNAP tag (3C cleavable) and a C-terminal double Strep II (DS) tag (TEV cleavable). Phosphomimetic mutant sequences were generated using the QuikChange Lightning Multi Site-Directed Mutagenesis kit (Agilent) according to the manufacturer’s instructions. Constructs were expressed in homemade *E. coli* BL21(DE3)-CodonPlus-RIL cells by induction with 0.4 mM IPTG and growth overnight at 18°C. 12 L of TB culture was generally required for sufficient sample for ITC experiments.

For Ndc80^N^ (and all variants) and Duo1^C-term^, cells were harvested by centrifugation and lysed by sonication in lysis buffer (50 mM Tris pH 8.0, 300 mM NaCl, 5 mM benzamidine, 1 mM β-mercaptoethanol, 5% glycerol) containing 10 µL benzonase nuclease (Merck), 1 mM PMSF and cOmplete EDTA-free protease inhibitor tablets (Roche). Lysate was clarified by centrifugation for 30 min at 38,000 x g and loaded onto 3 x 5 mL HiTrap TALON crude columns (Cytiva), washed with lysis buffer + 5 mM imidazole and eluted with lysis buffer + 250 mM imidazole. The eluate was loaded directly onto a 5 mL Strep-Tactin Superflow Plus cartridge (Qiagen) equilibrated in Strep buffer (20 mM Tris pH 8.0, 200 mM NaCl, 5 mM benzamidine, 1 mM β-mercaptoethanol, 5% glycerol). The column was washed with Strep buffer and eluted with Strep buffer + 3 mM desthiobiotin (IBA Lifesciences). Purification using affinity steps for both N- and C-terminal affinity tags allowed better yield of intact protein. TEV and 3C proteases were added to the Strep-Tactin eluate and incubated at 4°C overnight. Ndc80^N^ constructs containing the first 14 residues of Ndc80 bind weakly to TALON resin due to nearly tandem histidines at positions 10, 13 and 14. Therefore, the cleaved protein mixture was loaded onto a 3 x 5 mL HiTrap TALON crude column (Cytiva) equilibrated in reverse TALON buffer (20 mM Tris pH 8.0, 300 mM NaCl, 1 mM β-mercaptoethanol, 5% glycerol), washed with 1 CV reverse TALON buffer and the cleaved Ndc80^N^ construct eluted with a 4 CV gradient up to 30 mM imidazole. Cleaved Ndc80^N:40-70^ and Duo1^C-term^ do not bind TALON resin and thus were collected from the flow through. Desired fractions were pooled, concentrated by using a centrifugal filter and loaded onto a HiLoad 16/600 Superdex 75 pg (or 200 pg for Ndc80^N^) SEC column in SEC150 buffer (20 mM HEPES pH 7.5, 150 mM NaCl, 1 mM TCEP). Desired SEC fractions were concentrated, flash frozen in liquid nitrogen and stored at -80°C.

The full Ndc80^N^ does not survive freeze-thaw. For Ndc80^N^ ITC experiments, tags were not cleaved, thus the Strep-Tactin eluate was concentrated and dialysed into SEC150 buffer. A control ITC experiment showed no binding between the 6xHis-SNAP and DS tags and Bim1^EBH^.

### Cloning, expression and purification of Bim1^EBH^ and Bim1^EBH-mini^

Coding sequences from *S. cerevisiae* S288c for Bim1^EBH^ (Bim1 residues 171-290) and Bim1^EBH-mini^ (Bim1 residues 205-285) were cloned into a modified pETM11 vector containing an N-terminal 6xHis tag (TEV cleavable). Constructs were expressed in homemade *E. coli* BL21(DE3)-CodonPlus-RIL cells by growth at 37°C in autoinduction media (10 g/L tryptone, 5 g/L yeast extract, 0.5% glycerol, 0.05% glucose, 0.2% lactose, 25 mM (NH_4_)_2_SO_4_, 20 mM KH_2_PO_4_, 20 mM Na_2_HPO_4_) until OD600 of 0.6-0.8 was reached, followed by growth overnight at 18°C.

Cells expressing Bim1^EBH^ or Bim1^EBH-mini^ were harvested by centrifugation and lysed by sonication in lysis buffer (50 mM Tris pH 8.0, 300 mM NaCl, 5 mM benzamidine, 1 mM β-mercaptoethanol, 5% glycerol) containing 10 µL benzonase nuclease (Merck), 1 mM PMSF and cOmplete EDTA-free protease inhibitor tablets (Roche). Lysate was clarified by centrifugation for 30 min at 38,000 x g and loaded onto 5 x 5 mL HiTrap TALON crude columns (Cytiva) in lysis buffer, washed with lysis buffer + 5 mM imidazole and eluted with lysis buffer + 300 mM imidazole. TALON eluate was desalted into IEX buffer (20 mM Tris pH 8.0, 1 mM TCEP) + 100 mM NaCl using two tandem HiPrep 26/10 Desalting columns (Cytiva) and incubated with TEV protease overnight at 4°C or 20°C with rolling. This was loaded directly onto a 6 mL RESOURCE Q column (Cytiva), washed with IEX buffer + 100 mM NaCl and eluted over a 30 CV gradient to IEX buffer + 1 M NaCl. Desired fractions were concentrated using an Amicon Ultra 10 kDa MWCO centrifugal filter (Merck Millipore) and loaded onto a HiLoad 16/600 Superdex 200 pg (for Bim1^EBH^) or 75 pg (for Bim1^EBH-mini^) SEC column (Cytiva) equilibrated in SEC200 buffer. Desired were concentrated to > 1 mM, flash frozen in liquid nitrogen and stored at -80°C.

### Cloning, expression and purification of Ipl1-Sli15

Sli15 and Ipl1 coding regions were cloned from *S. cerevisiae* S288c genomic DNA into pU1 plasmid cassettes, with a double Strep II tag (TEV cleavable) fused onto the Sli15 N terminus (Zhang et al., 2016). High Five insect cells were infected by a P3 baculovirus made from DS-TEV-Sli15::Ipl1 in pU1 (Zhang et al., 2016). The cell pellet was lysed by sonication in lysis buffer (50 mM Tris pH 8.0, 500 mM NaCl, 3 mM benzamidine, 1 mM EDTA, 1 mM DTT, 5% glycerol) with cOmplete EDTA-free protease inhibitor tablets (Roche). The lysate was clarified by centrifugation and the supernatant loaded onto tandem Strep-Tactin Superflow Plus cartridges (Qiagen), washed with lysis buffer and eluted with lysis buffer + 3 mM desthiobiotin (IBA Lifesciences). The fractions containing the Sli15-Ipl1 complex were collected and loaded onto a HiLoad 16/600 Superdex 200 pg SEC column (Cytiva) equilibrated in 20 mM Tris pH 8.0, 300 mM NaCl, 1 mM EDTA, 1 mM DTT. The eluted Sli15-Ipl1 complex was concentrated using a centrifugal filter, flash frozen in liquid nitrogen and stored at -80°C.

### Preparation of phosphorylated Ndc80^N:1-70^

Phosphorylated Ndc80^N:1-70^(pNdc80^N:1-70^) was purified as described for Ndc80^N^ constructs except that 4 mM ATP, 10 mM MgCl_2_, 15 mM β-glycerophosphate and 10 mM NaF was added to the reverse TALON eluate following concentration. Ipl1-Sli15 kinase was then added to a 1:100 molar ratio of kinase to Ndc80^N:1-70^. The mixture was incubated for 16 h at 4°C, followed by addition of 20 mM EDTA to stop the phosphorylation reaction before loading of the sample onto SEC.

### Peptide synthesis

The SQIP (Ndc80^N:15-29^) (W-MDPHRFTSQIPTATS), mutant (Ndc80^N:15-29ΔSQIP^) (W-MDPHRFTASASTATS) and Ndc80^N-helix^ (W-NQGLTDMINKSIARTI) peptides were synthesised by Cambridge Research Biochemicals (now Biosynth) with N-terminal acetylation and C-terminal amidation. A tryptophan was included to measure peptide concentration using A_280_. Peptides were resuspended in 250 mM HEPES pH 7.5, 500 mM NaCl before dialysis into the appropriate buffer for subsequent use.

### Analytical size exclusion chromatography

Analytical SEC experiments were performed on an AktaMicro system (Cytiva) at 4°C in SEC150 buffer. Proteins were thawed on ice and centrifuged for 10 min at 4°C to remove aggregates. 50 µL reactions at 10 µM were set up and incubated at 20°C for 30 min before injecting onto the column. Agilent Bio SEC-5 LC 500 Å or 1000 Å (Agilent) or Superose 6 Increase 3.2/300 (Cytiva) columns were used as indicated. A_280_ chromatograms were plotted using GraphPad Prism and eluate fractions analysed by SDS-PAGE.

### SEC-MALS

SEC-MALS was performed with an Agilent 1200 series liquid chromatography system with an online DAWN HELEOS II system (Wyatt) equipped with a QELS+ module (Wyatt) and an Optilab rEX refractive index detector (Wyatt). 110 µL of protein sample was auto-injected onto a Superose 6 Increase 10/300 GL column (Cytiva) run at 0.5 mL/min in SEC150 buffer. Molecular weights were analysed using ASTRA (Wyatt) and data were plotted using GraphPad Prism.

### Isothermal titration calorimetry

ITC experiments were performed on a MicroCal PEAQ-ITC Automated calorimeter (Malvern Panalytical) at 20°C. Proteins for each experiment were dialysed overnight at 4°C or for 4 h at 20°C into the same batch of ITC buffer (20 mM HEPES pH 7.5, 150 mM NaCl, 1 mM TCEP). The cell was filled with 360 µL of sample. The pipette sample was titrated into the cell with an initial 0.5 µL injection followed by a further 19 injections of 2 µL sample, with a spacing of 180 s between injections and a stir speed of 750 RPM. Cell and pipette sample concentrations used for each experiment are specified in Results. All experiments were performed in triplicate with appropriate cell and pipette controls. Heat changes were integrated across the entire titration (excluding the first injection) and fitted with a single-site binding model using MicroCal PEAQ-ITC Analysis Software 1.52 (Malvern Panalytical). Stoichiometry was calculated assuming Bim1^EBH^ is a dimer.

### Protein structure prediction

Protein 3D structure prediction was performed with AlphaFold2 (Multimer v3) using a local implementation of ColabFold4 (Evans et al., 2022; Jumper et al., 2021; Mirdita et al., 2022). AlphaFold predictions were visualised in UCSF ChimeraX (Meng et al., 2023). Protein secondary structure prediction was performed using the PSIPRED workbench (Buchan & Jones, 2019; Jones, 1999).

### Multiple sequence alignment

Multiple sequence alignment was performed using Clustal Omega through the EMBL-EBI Job Dispatcher online tool (Madeira et al., 2024; Sievers & Higgins, 2021). Sequences were obtained from the UniProtKB database. Alignments were visualised in JalView 2.11.5.0 (Waterhouse et al., 2009), coloured by Clustal and conservation above 20% identity.

### NMR spectroscopy experiments

Uniformly ^15^N- or ^15^N, ^13^C-labelled proteins were expressed as described for unlabelled proteins but using M9 minimal media (6 g/L Na_2_HPO_4_, 3 g/L KH_2_PO_4_, 0.5 g/L NaCl) supplemented with 1.7 g/L yeast nitrogen base without NH_4_Cl and amino acids (YNB) (Sigma Y1251), 1 g/L ^15^NH_4_Cl and 4 g/L glucose (for ^15^N labelling) or 3.6 g/L ^13^C-glucose (for ^13^C, ^15^N labelling). For deuteration of nonlabile side chain protons, cells were grown successively on agar plates with 10%, 45% and 90% M9 minimal media with water replaced with 99% D_2_O (Sigma) supplemented with 1.7 g/L YNB, 1 g/L NH_4_Cl and 4 g/L glucose. Adapted colonies were used to inoculate 5 mL overnight cultures before growth of large-scale cultures in 99% D_2_O-M9 minimal media supplemented with 1.7 g/L YNB, 1 g/L ^15^NH_4_Cl and 3.6 g/L ^2^H,^13^C-glucose. Cells were harvested and the isotopically labelled protein purified as described for unlabelled protein.

NMR spectra were acquired on in-house Bruker Avance III spectrometers equipped with ^1^H detect TCI (800 MHz ^1^H resonance frequency) or ^13^C detect TXO (700 MHz ^1^H resonance frequency) cryoprobes. Spectra for the Bim1^EBH^ construct were acquired at the Biomolecular NMR Facility of the Francis Crick Institute (London, UK) on a Bruker Neo spectrometer with TCI cryoprobe (950 MHz ^1^H resonance frequency). All proteins were dialysed into fresh NMR buffer (20 mM HEPES pH 7.0, 100 mM NaCl, 1 mM TCEP) prior to experiments. Spectra for Ndc80^N^ and Bim1 constructs were collected at 278K using 25-50 µM sample and 298K using 100-200 µM sample, respectively. Data processing and analysis were carried out in Bruker Topspin 3.6, NMRFx 11.4.6 (Johnson, 2018; Koag et al., 2025) and Poky (Lee et al., 2021). The analysis of titration and dynamics experiments involved peak height analysis and fitting of exponential decays (T_2_) in Poky or NMRFx 11.4.6.

All 3D triple resonance experiments for backbone resonance assignments were acquired using nonuniform sampling (NUS), resulting in sparse data sets with 10-30% complex points. Data processing involved compressed sensing (CS) reconstruction with qMDD (Mayzel et al., 2014) or NMRView (Johnson, 2004). In general, the following 3D experiments (Bruker pulse sequence library) were acquired: HNCO, HN(CA)CO, HNCA, HN(CO)CA, HNCACB and CBCA(CO)NH (Cavanagh et al., 2007). Complex samples benefitted from additional HN(COCA)NNH experiments. Backbone assignments were based on correlations of HN, N, Cα, Cβ and C’ chemical shifts in Poky supported with automatic chemical shift assignments in MARS (Jung & Zweckstetter, 2004) and curated manually using in-house scripts.

For Bim1 constructs, 3D data sets employed transverse relaxation optimised spectroscopy (TROSY)-based 3D triple resonance experiments with side chain deuterium decoupling in the case of ^2^H,^13^C,^15^N-labelled samples and INEPT based 3D experiments for ^13^C,^15^N-labelled samples (Morris, 1980; Pervushin, 2000). For Ndc80 constructs, either BEST (Band-selective Excitation Short Transient)-TROSY or conventional INEPT pulse sequences were employed. To complete the assignment of backbone imino resonances of prolines and for residues with rapid solvent exchange, additional ^15^N, ^13^C correlations using ^1^H start versions of ^13^C-detected CON, CACON and CANCO experiments were obtained (Bruker pulse sequence library).

Most titration experiments were recorded as 2D FHSQC data sets. Relative peak intensities were normalised to non-binding residues located at or near the N-terminus of the constructs and expressed as I/I_0_, where I and I_0_ correspond to the peak intensities of a given residue in the bound and free states, respectively. ^15^N transverse relaxation (T_2_) experiments were acquired as interleaved, temperature compensated pseudo-3D experiments with multiple relaxation delays (e.g. 8.48 (repeated twice), 16.96, 33.92, 50.88, 67.84, 101.76, 135.68, 169.6, 203.52,685 237.44, 271.36 ms). Secondary chemical shifts were calculated based on differences between experimental Cα and Cβ chemical shifts and calculated random coil values accounting for experimental temperature, salt concentration, pH and residue phosphorylation state (if applicable) (Hendus-Altenburger et al., 2019; Kjaergaard et al., 2011; Kjaergaard & Poulsen, 2011). Weighted CSPs were calculated as √(Δδ^1^H)^2^ + (0.2(Δδ^15^N)^2^), with Δδ^1^H and Δδ^15^N being the chemical shift differences between free and bound states in the proton and nitrogen dimensions, respectively.

### NMR phosphorylation experiments

For on-magnet phosphorylation, a 2D ^1^H, ^15^N FHSQC spectrum for Ndc80^N:1-70^ was first acquired in NMR phosphorylation buffer (20 mM HEPES pH 7.0, 100 mM NaCl, 1 mM TCEP, 2 mM ATP, 5 mM MgCl_2_, 15 mM β-glycerophosphate, 10 mM NaF) as described above for Ndc80 samples before addition of Ipl1-Sli15 at a 1:5000 ratio of kinase to Ndc80^N:1-70^. Spectra were acquired approximately every 10 minutes to monitor changes in the spectra due to protein phosphorylation. For pNdc80^N:1-70^ dynamics and titration experiments, phosphorylation was quenched by addition of 10 mM EDTA (from a 0.5 M EDTA solution, pH 9.0) at ∼80% completion of Ser37 phosphorylation and ∼35% completion of Thr54 phosphorylation. Only signals from the phosphorylated species were considered in analysis.

### NMR data visualisation

NMR spectra and bar charts from titration and dynamics experiments were output from the appropriate software as .ps files which were opened in Adobe Illustrator, in which data were format to facilitate data visualisation. I/I_0_ values from titration experiments and R_2_ rates from transverse relaxation experiments were converted into a B-factor-style file format using in-house scripts and plot onto the relevant AlphaFold2 predictions as B-factors using PyMOL 2.5.0 (Schrodinger) run with the loadBfacts.py script (Gatti-Lafranconi, 2014).

### Crosslinking mass spectrometry

Protein crosslinking reactions were carried out with 2 mM sulfo-SDA or BS3 as specified. Sulfo-SDA and complex were mixed and incubated on ice for 15 min before being crosslinked for 15 s with 365 nm UV radiation from a home build UV LED setup. BS3 and complex were mixed and incubated on ice for 30 min. Crosslinking reactions were quenched by addition of 50 mM Tris. The quenched solution was reduced with 5 mM DTT and alkylated with 20 mM iodoacetamide. SP3 protocol as described in (Batth et al., 2019; Hughes et al., 2019) was used to clean-up and buffer exchange the reduced and alkylated protein. Shortly, proteins were washed with ethanol using magnetic beads for protein capture and binding. The proteins were resuspended in 100 mM NH_4_HCO_3_ and were digested with trypsin (Promega) at an enzyme-to-substrate ratio of 1:20, and Rapigest 0.1% (Waters). Digestion was carried out overnight at 37 °C. Clean-up of peptide digests was carried out with HyperSep SpinTip P-20 (ThermoFisher Scientific) C18 columns, using 60% Acetonitrile as the elution solvent. Peptides were then evaporated to dryness using Speed Vac Plus (Savant).

Dried peptides were resuspended in 30% acetonitrile and were fractionated by size exclusion chromatography using a Superdex 30 Increase 3.2/300 column (Cytiva) at a flow rate of 20 µL/min using 30% (v/v) acetonitrile 0.1 % (v/v) trifluoroacetic acid as a mobile phase. Fractions were taken every 5 min, and fractions 2-7 containing crosslinked peptides were collected. Dried peptides were suspended in 3% (v/v) acetonitrile and 0.1 % (v/v) formic acid and analysed by nanoscale capillary LC-MS/MS using a Vanquish Neo HPLC (ThermoFisher Scientific) to deliver a flow of 300 nL/min. Peptides were trapped on a C18 Acclaim PepMap100 5 μm, 0.3 μm x 5 mm cartridge (ThermoFisher Scientific) before separation on Aurora Ultimate C18, 1.7 μm, 75 μm x 25 cm (Ionopticks). Peptides were eluted on optimised gradients of 90 min and interfaced using a nanoFlex ionisation source to a tribrid quadrupole Orbitrap mass spectrometer (Orbitrap Ascend, ThermoFisher Scientific). Mass spectrometry data were acquired in data dependent mode with a TopS method with 3 s cycle times, high resolution scans full mass scans were carried out (R = 120,000, *m/z* 400 – 1550) followed by higher energy collision dissociation with stepped collision energy range 21, 30, 34 % normalised collision energy. The tandem mass spectra were recorded (R=60,000, isolation window *m/z* 1, dynamic exclusion 50 s).

### Crosslinking mass spectrometry data analysis

Xcalibur raw files were converted to MGF files using ProteoWizard (Chambers et al., 2012) and crosslinks were analysed using xiSEARCH (Mendes et al., 2019). Search conditions used 2 maximum missed cleavages with a minimum peptide length of 5. Variable modifications used were carbmidomethylation of cysteine (57.02146 Da) and methionine oxidation (15.99491 Da). False discovery rate was set to 1%.

### Intact protein mass spectrometry

Protein samples were analysed by LC-MS using a Vanquish HPLC (ThermoFisher Scientific) with mobile phase A – 90% water, 9.9% acetonitrile with 0.1% formic acid (FA) and mobile phase B – 80% acetonitrile, 19.9% water, and 0.1% FA. Delivered flow was 300 µL/min. Proteins were trapped on a MAbPac RP 2.1 x10 mm trapping column for 2 min before separation using a MAbPac RP 2.1 x 50 mm liquid chromatography column. The gradient was from 10% to 30% B over 2 min before increasing to 55% B over the next 8 min at 80°C. Proteins were eluted and ionised with a HESI ion source onto an orbitrap mass spectrometry (Exploris MX, ThermoFisher Scientific). Mass spectrometry data were acquired in in a broadband detection mode (R = 120,000, *m/z* 600 – 2000, intact protein mode) in positive ion mode (AGC = 100%, µscans = 10, in source energy = 10 eV). Protein deconvolution was carried out using a sliding window Xtract (isotopically resolved) deconvolution to generate monoisotopic protein masses.

### Yeast strain construction

Yeast strains constructed for this study are listed in Table S2. All Ndc80 mutant rescue strains were constructed from the Ndc80-AID strain from Kyle Muir (Muir et al., 2023). The *NDC80* gene with the appropriate mutation and a C-terminal 3xHA tag was cloned into a pRS304 vector for integration at the exogenous *TRP1* locus (Sikorski & Hieter, 1989). 500 bp of the *S. cerevisiae* strain S288c genomic DNA sequence upstream and downstream of the endogenous *NDC80* locus was included in the rescue vectors 5’ and 3’ to the rescue *NDC80* sequence, respectively, such that the rescue Ndc80 gene product was regulated by endogenous Ndc80 promoter and terminator regions. All rescue vectors were linearised using BsgI (NEB) to form the transformation cassette. 1 µg of the appropriate transformation cassette was transformed into Ndc80-AID cells from exponential culture using the lithium acetate method as described (Gietz & Schiestl, 2007). Cells were heat shocked for 45 min at 42°C. Transformants were plated on tryptophan dropout agar (-TRP) plates and grown at 30°C for 3 days. Transformant colonies were re-streaked onto-TRP plates and grown at 30°C for 3 days to ensure robust selection. Colonies were then screened for integration of the transformation cassette by analytic PCR using extracted genomic DNA as the template and primers amplifying across the 5’ and 3’ integration junctions. Single colonies were streaked into 1 cm^2^ patches on-TRP plates and grown at 30°C for 3 days. Cells from the patch were resuspended in 100 µL 200 mM lithium acetate 1% SDS solution and the genomic DNA was extracted as outlined in (Lõoke et al., 2011), except that precipitated DNA was resuspended in 30 µL Milli-Q water. Analytic PCR products were analysed by agarose gel electrophoresis. Exogenous expression of the Ndc80 mutants was confirmed by Western blotting and further validated by sequencing.

### Western blotting

To assess ectopic Ndc80 expression from constructed yeast strains, 1 mL of yeast cells at OD600 of 1 were harvested by centrifugation at 5,000 x g for 5 min. Cells were lysed with the post-alkaline method as in (Kushnirov, 2000), but using 50 µL 1X NuPAGE LDS Sample Buffer (Invitrogen) supplemented with DTT instead of SDS sample buffer. 1-10 µL of sample was loaded onto SDS-PAGE using a 4-12% NuPAGE Bis-Tris gel (Invitrogen) run at 200 V for 40 min. Proteins were transferred from the SDS-PAGE gel to a nitrocellulose membrane using a Trans-Blot Turbo Mini 0.2 µm Nitrocellulose Transfer Pack (Bio-Rad) and Trans-Blot Turbo Transfer System (Bio-Rad) following the manufacturer’s protocol. Membranes were blocked by incubation by rolling with 10 mL phosphate-buffered saline with 0.1% TWEEN 20 (Sigma-Aldrich) (PBS-T) + 5% milk powder for 30 min at 20°C. To detect Ndc80-HA_3_, we used rabbit anti-HA (Sigma-Aldrich H6908, 1:1000 dilution). Rat anti α-tubulin (Bio-Rad MCA78G, 1:3000 dilution) was used as the loading control. Primary antibody was added directly to the milk suspension, and membranes were incubated by rolling for 1 h at 20°C or overnight at 4°C then washed thoroughly with PBS-T (3 x 10 min with rolling at 20°C). HRP-conjugated anti-rabbit (Invitrogen SA1-200, 1:3000 dilution) or anti-rat (Santa Cruz Biotechnology sc-2302, 1:3000 dilution) secondary antibody was added as appropriate to 10 mL PBS-T and the membrane incubated for 1 h at 20°C. The membrane was washed thoroughly with PBS-T and blots visualised using Amersham ECL Prime detection reagent (Cytiva) and a ChemiDoc imaging system (Bio-Rad).

### Auxin depletion assays

Single colonies for each strain were used to inoculate 10 mL overnight cultures in YEPD (30°C, 220 RPM). The OD600 of each culture was measured in triplicate, cells were diluted to OD600 = 1 and serially diluted 1:9 (vol/vol) over four further dilutions. 4 µL of each dilution was spot onto prewarmed YEPD, YEPD + 0.5 mM IAA, YEPD + 15 µg/mL benomyl or YEPD + 15 µg/mL benomyl + 0.5 mM IAA plates as appropriate and grown for 3 days at 30°C before imaging.

### Optical trap rupture force assay

All rupture force experiments were performed on an in-house LUMICKS C-Trap Edge setup. The procedure followed was very similar to that published by colleagues (Muir et al., 2023). SNAP-tagged Ndc80c or Ndc80c mutant was biotinylated using SNAP-biotin (NEB) and linked to 0.9 µm-diameter streptavidin-coated polystyrene beads (LUMICKS). 3.5 pM beads were incubated with 30 nM Ndc80c for 1 h at 22°C and washed twice in PBS.

Glass 22x22 mm coverslips (NEXTERION, Schott) were incubated for at least 48 h at 22°C in a 1:10 (w/w) solution of mPEG-Silane (30 kDa, PSB-2014, Creative PEGWorks) and Biotin-PEG-Silane (3.4kDa, Laysan Bio Inc.) in 96 % (v/v) ethanol and 0.2 % (v/v) HCl. The coverslips were then washed in ethanol and ultrapure water, dried with an air duster and assembled into a flow cell on a slide using double-sided tape (Adhesive Research AR-90880 precisely cut with a Graphtec CE6000 cutting plotter). The chamber was first perfused with one chamber volume of BRB80 (80 mM PIPES pH 6.9, 2 mM MgCl_2_, 0.5 mM EGTA, 1 mM DTT), then with one chamber volume of 5% Pluronic-F127 in BRB80 and immediately flushed out with 8 chamber volumes of BRB80. One chamber volume of Neutravidin (25 µg/mL) diluted in BRB80 was left to incubate for 5 min at 22°C and then flushed out with 4 chamber volumes of BRB80. Biotin-GMPCPP stabilised microtubule seeds (20% HiLyte 488-tubulin, 20% biotin-tubulin 60% tubulin, from Cytoskeleton, Inc.) were then injected and let to adhere to the neutravidin for 5 min. The chamber was then washed with 10 chamber volumes of BRB80. Ndc80c coated beads were added to the reaction mixture (1x BRB80, 16.7 μM tubulin, 80 μM GTP, 1 mg/mL BSA). Bim1 was added to a final concentration of 70 nM and Dam1 was added to a final concentration of 30 nM where appropriate. The microscope stage was kept at 28°C with an objective and condenser heater (LUMICKS). 488nm HiLyte tubulin seeds were briefly localised in TIRF mode. Microtubule growth, bead capture and rupture experiments were carried out using interference reflection microscopy (IRM). Data were collected for a maximum of 90 min after addition of the reaction mixture.

Beads were held in the laser trap approximately 200 nm above the surface and the position was continually monitored at 40 KHz. Trap stiffness was calibrated using a power spectrum method and was in the range of 0.025-0.035 pN/nm. Beads were allowed to bind to the matrix near to the tip of the microtubule. The laser-trap was moved at 10 μm/s until the bead ruptured from the tip. Data were recorded using Bluelake software (LUMICKS) and rupture forces were manually determined by examining force and position versus time traces using PRISM. 95% confidence intervals (CIs) were generated for the median rupture forces. Normality test showed a non-parametric analysis should be performed, therefore statistical significance for the difference in medians between each sample was determined using a Kruksal-Wallis test.

### Microtubule tip tracking assay

All microtubule tip tracking experiments were performed on an in-house LUMICKS C-Trap Edge setup. Coverslips coated with biotin-GMPCPP stabilised microtubule seeds were prepared as described for rupture force experiments but using imaging buffer (BRB80 + 150 mM KCl, 0.1 mg/mL K-Casein, 0.1 mg/mL BSA, 40 µM DTT, 64 mM D-glucose, 160 µg/mL glucose oxidase, 20 µg/mL catalase and 0.2% methylcellulose). Ndc80c-SNAP was labelled using SNAP-Surface Alexa Fluor 647 (NEB). A reaction mixture containing imaging buffer, 16.7 μM tubulin, 80 μM GTP and 7 nM Ndc80c-SNAP-647 with or without 70 nM Bim1 (unlabelled) was added to the chamber. 150 mM KCl and a low concentration of Ndc80c was used to prevent Ndc80c binding to the microtubule lattice. The microscope stage was kept at 28°C with an objective and condenser heater (LUMICKS). Fluorescence image sequences were acquired for 3–5 min at 1 frame s⁻¹ using a 639 nm laser at 2% power with an exposure time of 250 ms. IRM images were acquired simultaneously at 5 frames s⁻¹ with an exposure time of 10 ms and subsequently averaged to generate a final frame rate of 1 frame s⁻¹. Data acquisition was continued for a maximum of 90 min following addition of the reaction mixture.

Images were processed using Fiji (version 2.18.0) (Schindelin et al., 2012). Figures were assembled in Adobe Illustrator 2024. IRM and TIRF movies were manually aligned using microtubule seed and crossover locations, the background was subtracted from each channel, and the channels merged. Microtubules were manually traced as straight lines and kymographs were created using the ReSlice function.

## Data availability

NMR backbone assignments were deposited to the BMRB database (https://bmrb.io) with the following accession numbers: Ndc80^N:1-70^, 53896; pNdc80^N:1-70^, 53909; Ndc80^N^, 53897; Bim1^EBH^, 53900; Bim1^EBH-mini^, 53899. The mass spectrometry proteomics data have been deposited to the ProteomeXchange Consortium (http://proteomecentral.proteomexchange.org/) via the PRIDE partner repository (Perez-Riverol et al., 2025) with the dataset identifier PXD082987.

## Author Contributions

Yvonne B. Winterborn: conceptualization, data curation, formal analysis, investigation, methodology, validation, visualization, writing – original draft, review & editing. Christopher Batters: investigation, formal analysis, methodology, writing – review & editing. Tomos E. Morgan: investigation, formal analysis. Stefan M.V. Freund: conceptualization, data curation, formal analysis, investigation, validation, writing – review & editing. David Barford: conceptualization, formal analysis, funding acquisition, methodology, project administration, resources, supervision, validation, writing – review & editing.

## Acknowledgements

We are grateful to C.H.W. Yu for initial NMR investigations on this project; to K. Muir for yeast strains and discussions; to Z. Zhang for providing Ipl1-Sli15 and Dam1c, and J. Yang for providing some Ndc80c; to K. Turton for help with insect cell expression; to S.H. McLaughlin for help with biophysics; and to J. Grimmett, T. Darling, and I. Clayson for scientific computing. This work was supported by the Francis Crick Institute through provision of access to the MRC Biomedical NMR Centre. The Francis Crick Institute receives its core funding from Cancer Research UK (CC1078), the UK Medical Research Council (CC1078), and the Wellcome Trust (CC1078).

---

**Figure S1.**
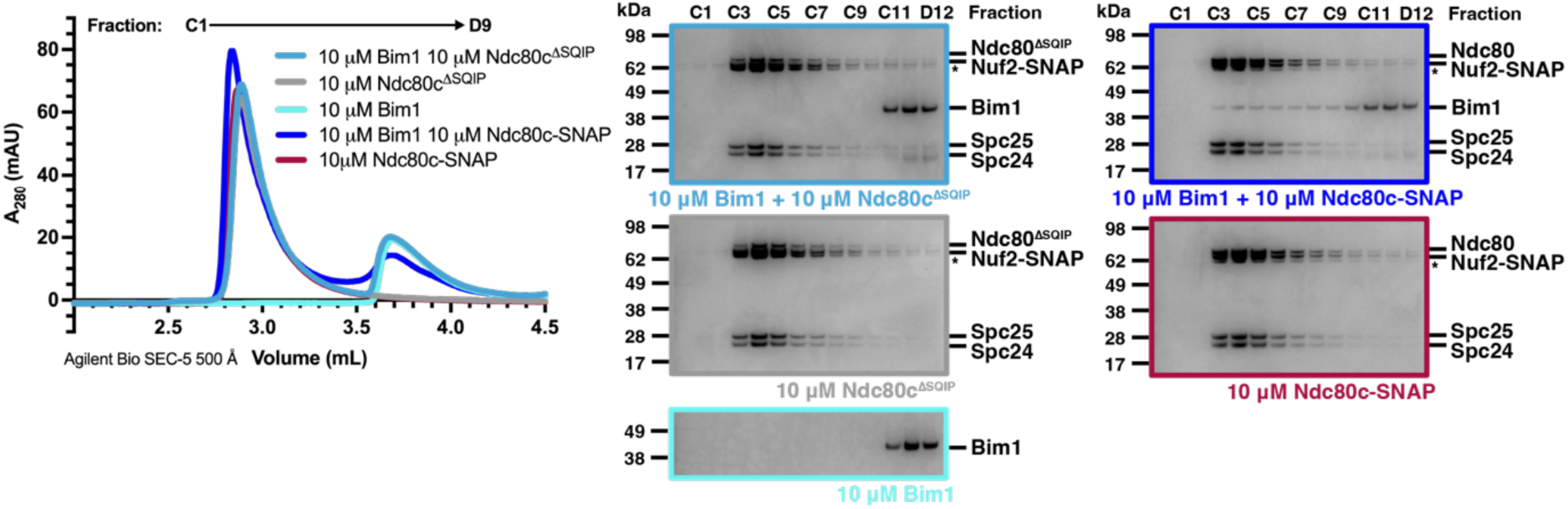
– The Ndc80^N^ SQIP motif is required for stable Bim1-Ndc80c interaction as assessed by analytic SEC. Chromatogram (left) and corresponding SDS-PAGE analysis (right) of eluate fractions of Bim1 and Ndc80c-SNAP or Ndc80c^ΔSQIP^-SNAP injected onto an Agilent Bio SEC-5 500 Å column individually and together.

**Figure S2.**
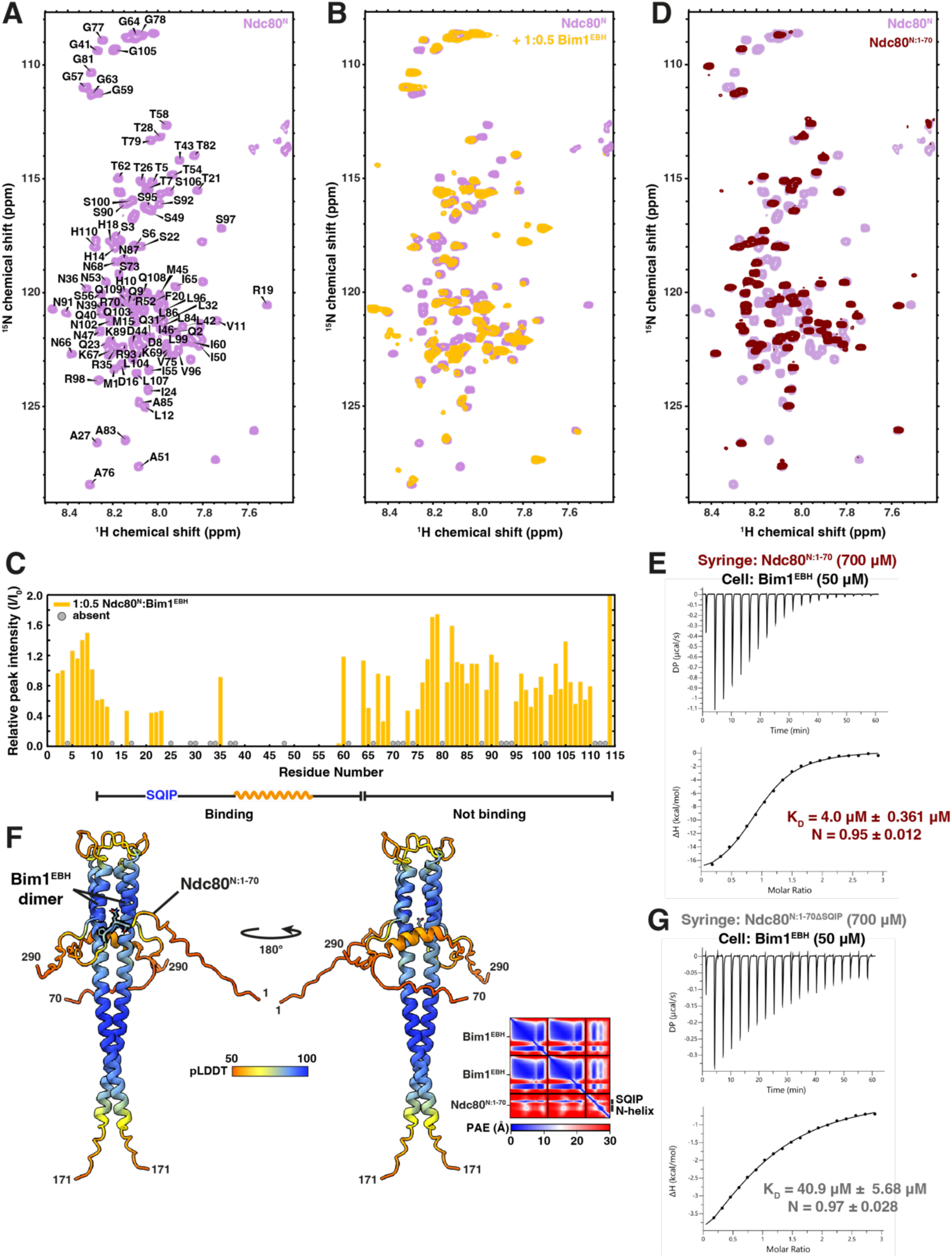
– NMR investigations into Ndc80^N^ and Ndc80^N:1-70^. **(A)** 2D ^1^H, ^15^N HSQC spectrum of ^15^N-labelled Ndc80^N^, with partial backbone assignments. 23 residues are unassigned. **(B)** Overlay of 2D ^1^H, ^15^N HSQC spectra of ^15^N-labelled Ndc80^N^ alone and with a 1:0.5 ratio of Bim1^EBH^. **(C)** Relative peak intensities (I/I0) of the 2D ^1^H, ^15^N HSQC spectrum of Ndc80^N^ with 1:0.5 ratio of Bim1^EBH^. Peak attenuation indicates that residues 10-63 are generally involved in binding Bim1^EBH^ and residues 1-9 and 64-114 are not. Peak intensities are normalised relative to Ser3. ‘Absent’ indicates peaks that are unassigned in the 2D ^1^H ^15^N HSQC spectra of free Ndc80^N^ including prolines which lack amide resonances. **(D)** Overlay of 2D ^1^H, ^15^N HSQC spectra of ^15^N-labelled Ndc80^N^ and ^15^N-labelled Ndc80^N:1-70^. **(E)** ITC of Ndc80^N:1-70^ with Bim1^EBH^. **(F)** Bim1^EBH^ and Ndc80^N:1-70^ AlphaFold2 prediction coloured by pLDDT score (left). PAE plot for the prediction (right). **(G)** ITC of Ndc80^N:1-70ΔSQIP^ with Bim1^EBH^.

**Figure S3.**
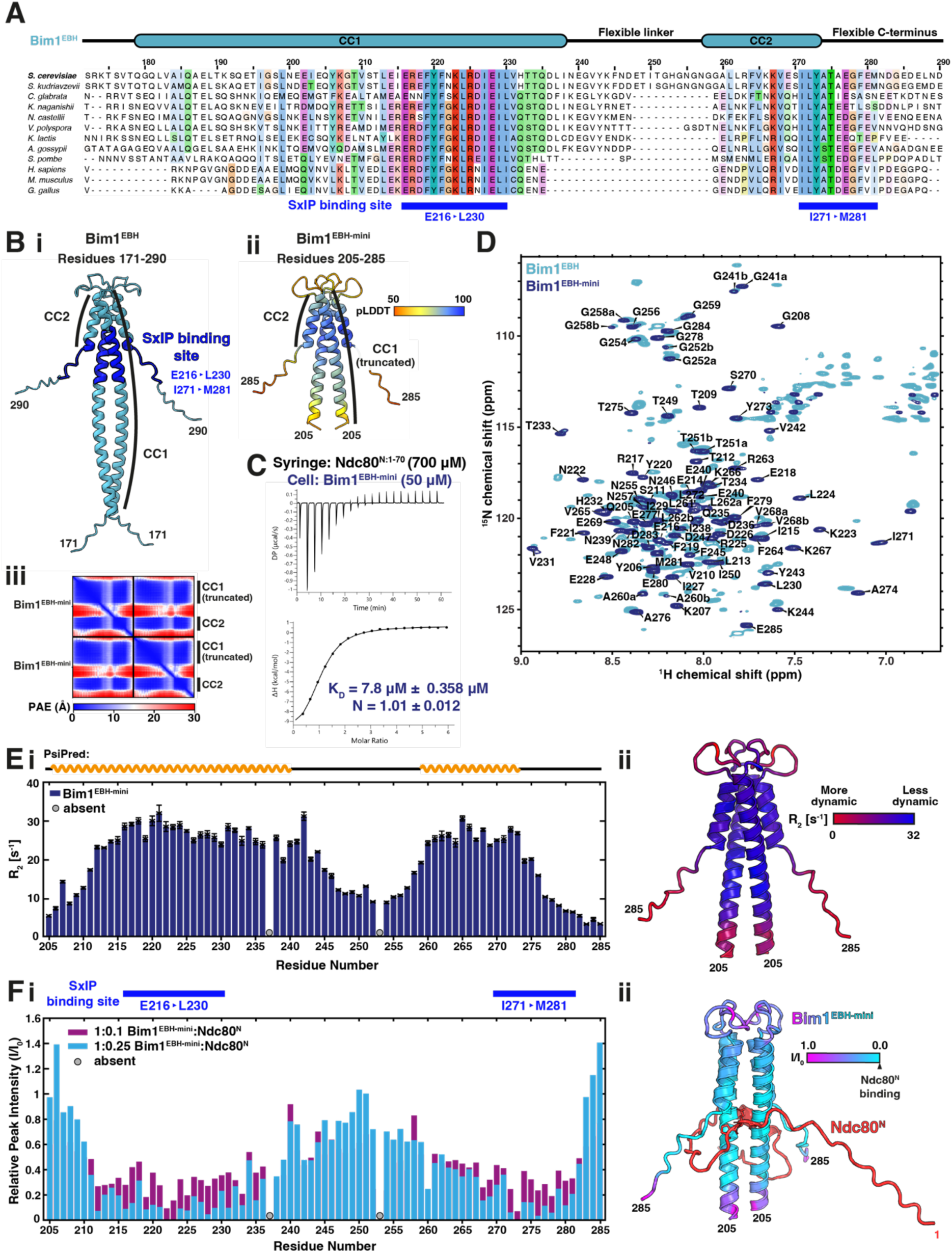
– NMR investigations into the Ndc80-binding of Bim1. **(A)** MSA across eukaryotes of the EBH domain from EB-family proteins, with the secondary structure of the Bim EBH domain shown above. The EBH domain is a coiled-coil dimer; there is a break in the coiled-coil between CC1 and CC2, which folds back to form a four-helical bundle at the C-terminus. The flexible linker between CC1 and CC2 is extended in *Saccharomyces* species. The SxIP motif binding site includes residues from CC1 (Glu216-Leu230), CC2 and the flexible C-terminus (residues Ile271-Met281) at the base of the four-helical bundle. **(B)(i)** Bim1^EBH^ AlphaFold2 prediction showing the SxIP motif binding site. **(ii)** AlphaFold2 prediction of Bim1^EBH-mini^, residues 205-285, coloured by pLDDT. The Bim1^EBH-mini^ construct comprises the SxIP motif binding site but with a truncated CC1. **(iii)** PAE plot for the Bim1^EBH-mini^ AlphaFold2 prediction. **(C)** ITC of Ndc80^N:1-70^ with Bim1^EBH-mini^. **(D)** Overlay of 2D ^1^H, ^15^N HSQC spectra of perdeuterated Bim1^EBH^ and ^15^N-labelled Bim1^EBH-mini^ with Bim1^EBH-mini^ backbone assignments labelled. Cross peaks present in both constructs overlay well, indicating that the Bim1^EBH-mini^ secondary structure elements are retained in Bim1^EBH^. **(E)(i)** R2 (1/T2) relaxation rates of ^15^N-labelled Bim1^EBH-mini^. Higher R2 rates are observed for residues which are less dynamic (folded conformation), whereas lower R2 rates indicate more dynamic residues (disordered). The higher R2 rates correspond well to helices predicted by PsiPred. Error bars are the likely error based on Gaussian noise from peak heights. **(ii)** R2 rates from (i) plotted onto the AlphaFold2 prediction of Bim1^EBH-mini^. Lower R2 rates at the N-terminus of the sequence suggest some fraying of the truncated CC1. **(F)** Relative peak intensities (I/I0) of the 2D ^1^H, ^15^N HSQC spectra of ^15^N-labelled Bim1^EBH-mini^ with a 1:0.1 and 1:0.25 ratio of Ndc80^N^. Peaks are normalised relative to Ile250. ‘Absent’ indicates peaks that are not assigned in the ^1^H, ^15^N 2D HSQC spectra of free Bim1^EBH-mini^ including prolines which lack amide resonances. **(G)** Relative peak intensities (I/I0) of 1:0.25 Bim1^EBH-mini^: Ndc80^N^ plotted on the AlphaFold2 prediction of the interaction. Bim1^EBH-mini^ is coloured by I/I0.

**Figure S4.**
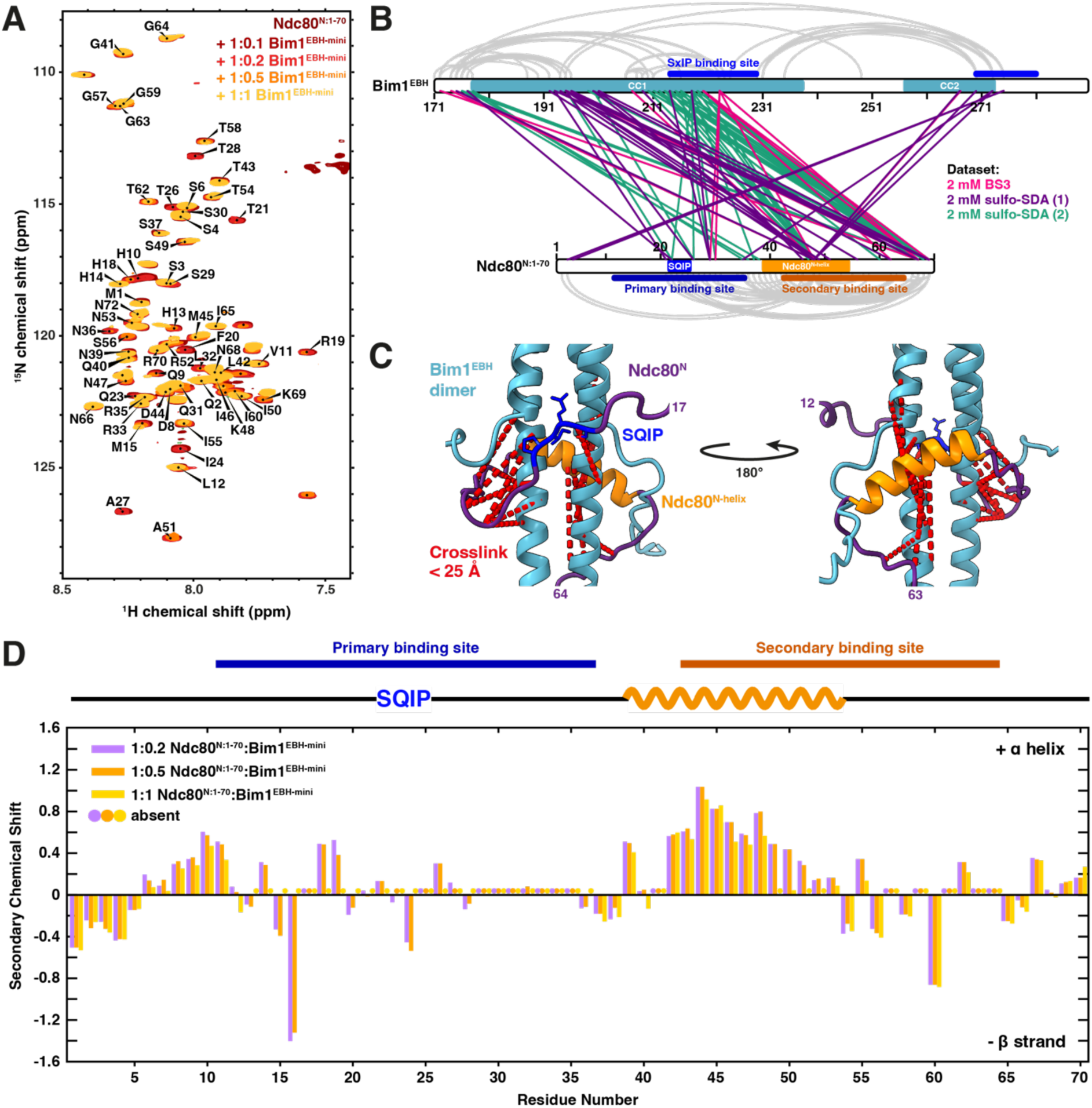
– NMR investigations of Ndc80(1-70) in complex with Bim1^EBH-mini^. **(A)** Overlay of 2D ^1^H, ^15^N HSQC spectra of Ndc80^N:1-70^ titrated with indicated concentrations of Bim1^EBH-mini^. **(B)** Map of crosslinks observed for a Bim1^EBH^:Ndc80^N:1-70^ complex crosslinked with 2 mM sulfo-SDA or 2 mM BS3. **(C)** Selected crosslinks from (B) plotted onto the Bim1^EBH^:Ndc80^N^ AlphaFold2 prediction. Crosslinks shown are between residues that are near the Bim1^EBH^-SxIP binding site and that satisfy the physical distance constraint of sulfo-SDA (25 Å). **(D)** Secondary chemical shifts corresponding to secondary structure propensity of ^13^C,^15^N-labelled Ndc80^N:1-70^ in complex with increasing concentrations of Bim1^EBH-mini^ (three bars for each residue). Many peaks are absent due to peak attenuation in the presence of Bim1^EBH-mini^. Present peaks indicate that secondary structure propensity of Ndc80^N:1-70^ does not change significantly upon binding.

**Figure S5.**
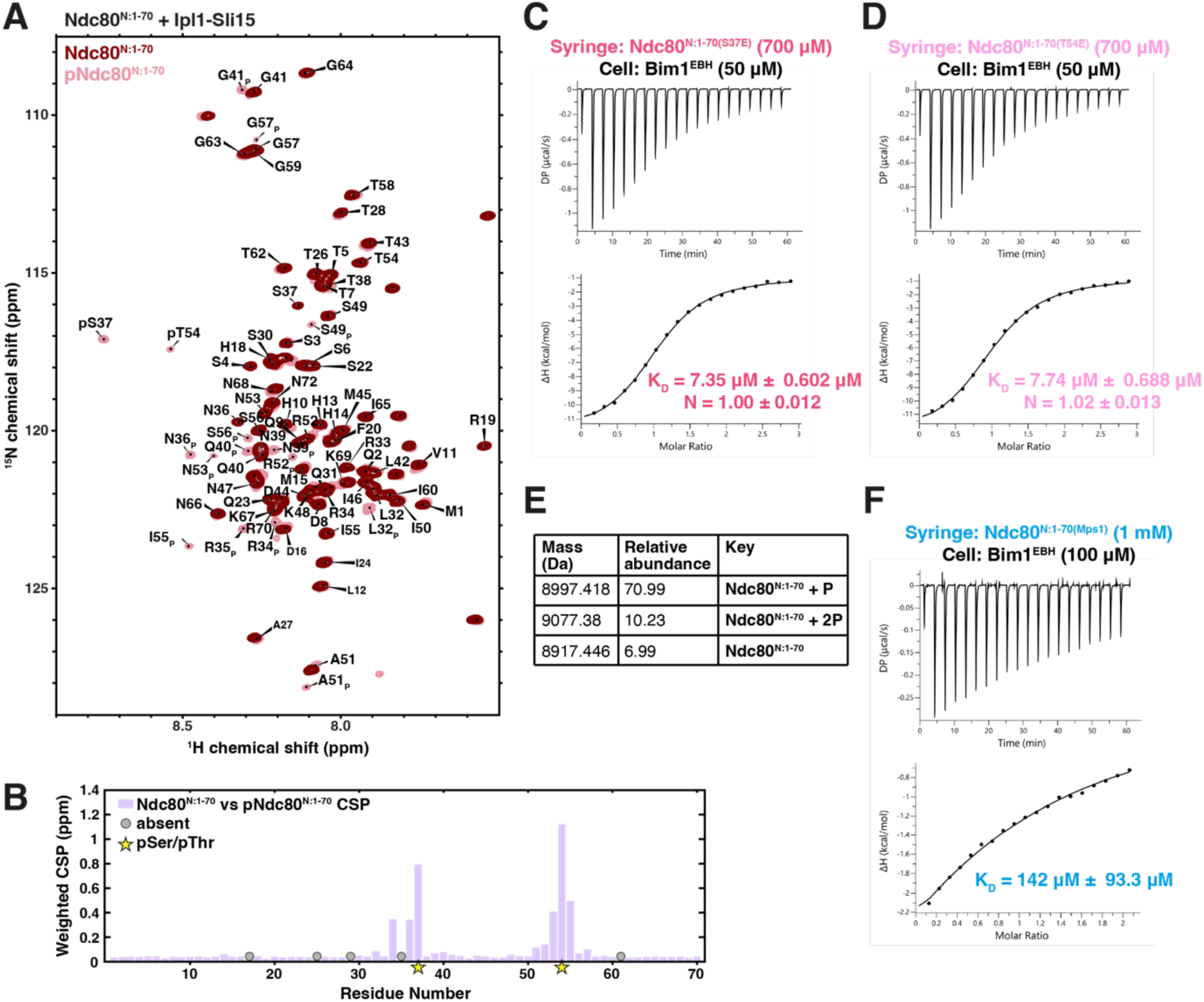
– Additional investigation into phosphorylation of the Ndc80 N-tail. **(A)** Overlay of 2D ^1^H, ^15^N HSQC spectra of ^13^C,^15^N-labelled pNdc80^N:1-70^ and at completion of phosphorylation with Ipl1-Sli15, pNdc80^N:1-70^, with pNdc80^N:1-70^ backbone assignments labelled. Ser37 and Thr54 are phosphorylated by Ipl1 (pS37 and pT54). XP denotes the new position of residue X affected by Ndc80^N:1-70^ phosphorylation. **(B)** Weighted chemical shift perturbations (CSPs) of the peak changes seen in (A). Residues close to pSer37 and pThr54 experience large CSPs, consistent with the chemical environment change upon phosphorylation of nearby residues. **(C)** ITC of Ndc80^N:1-70(S37E)^ with Bim1^EBH^. **(D)** ITC of Ndc80^N:1-70(T54E)^ with Bim1^EBH^. **(E)** Intact mass spectrometry analysis of pNdc80^N:1-70^ used for ITC experiments. **(F)** ITC of Ndc80^N:1-70(Mps1)^ (Ndc80^N:1-70^ with all Mps1 sites within the Bim1-binding mutated to glutamate: T21E S22E S37E T38E T43E) with Bim1^EBH^. Stoichiometry could not be accurately determined due to low c-value.

**Figure S6.**
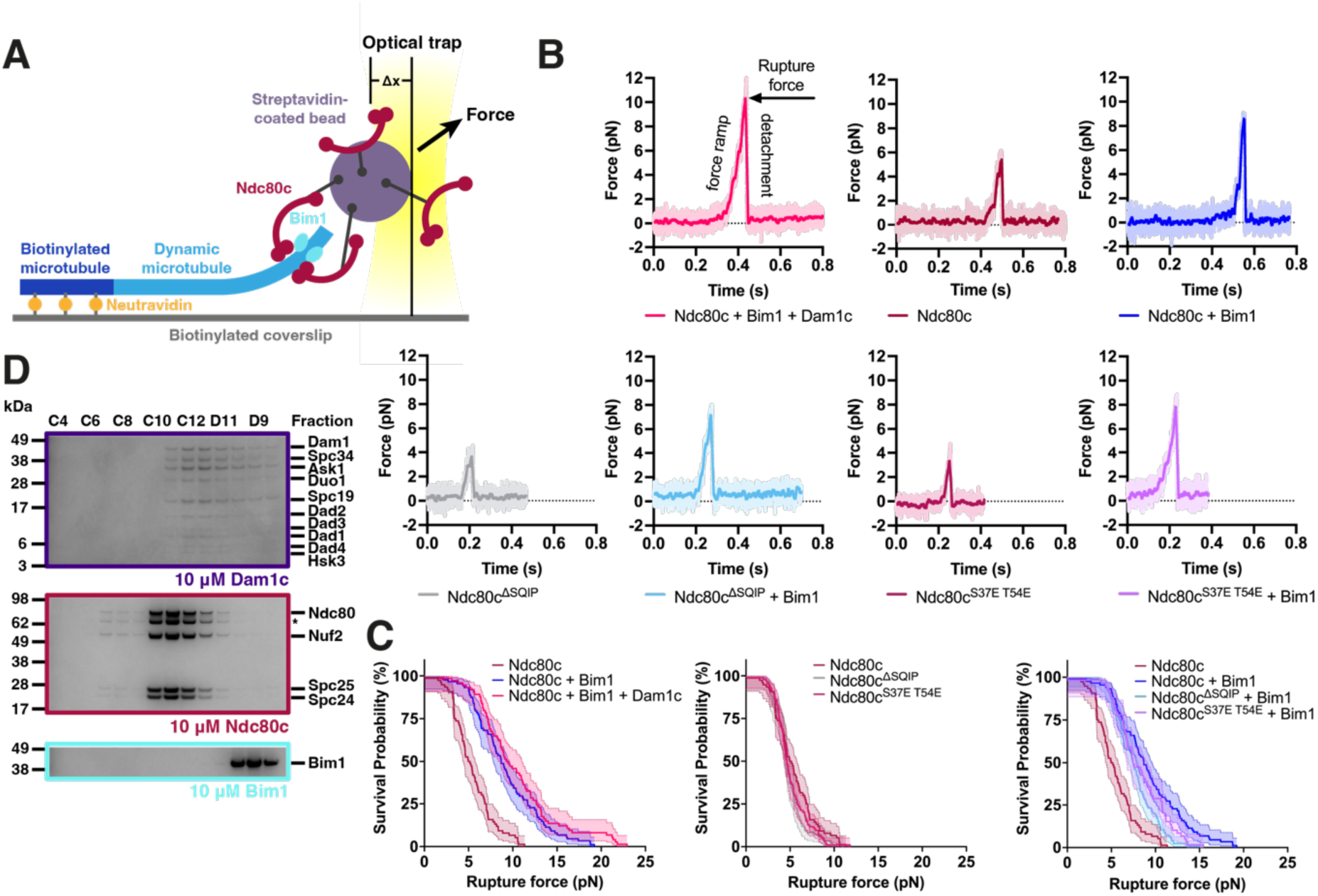
– Additional details for optical trap experiments and Bim1-Ndc80c-Dam1c analytical SEC. **(A)** Schematic summarising the optical trap rupture force assay. Ndc80c or Ndc80c mutants were attached to streptavidin-coated beads via a SNAP-biotin tag, and a dynamic microtubule was grown from a biotinylated microtubule seed immobilised onto a biotinylated coverslip with neutravidin. Beads held in the optical trap laser were brought close to microtubules to allow Ndc80c-microtubule attachments to form then withdrawn to determine the rupture force of the attachment. Experiments were performed with and without Bim1 and Dam1c in the flow cell. Force = kΔx (k = optical trap stiffness). **(B)** Example traces showing force detachment for each dataset collected. A trace with rupture force close to the median for the dataset is shown. Light dots represent raw data, and the solid line shows the same data after smoothing with a 20 ms sliding boxcar average. Rupture force is at the peak of the trace. **(C)** Rupture force survival probability plots displayed as Kaplan-Meier survival curves for Ndc80c with Bim1 and Bim1 and Dam1c (left), Ndc80c mutants (middle) and Ndc80c mutants with Bim1 (right). **(D)** SDS-PAGE analysis of Dam1c, Ndc80c and Bim1 eluate fractions from the analytical SEC shown in Figure 8C.

**Figure S7.**
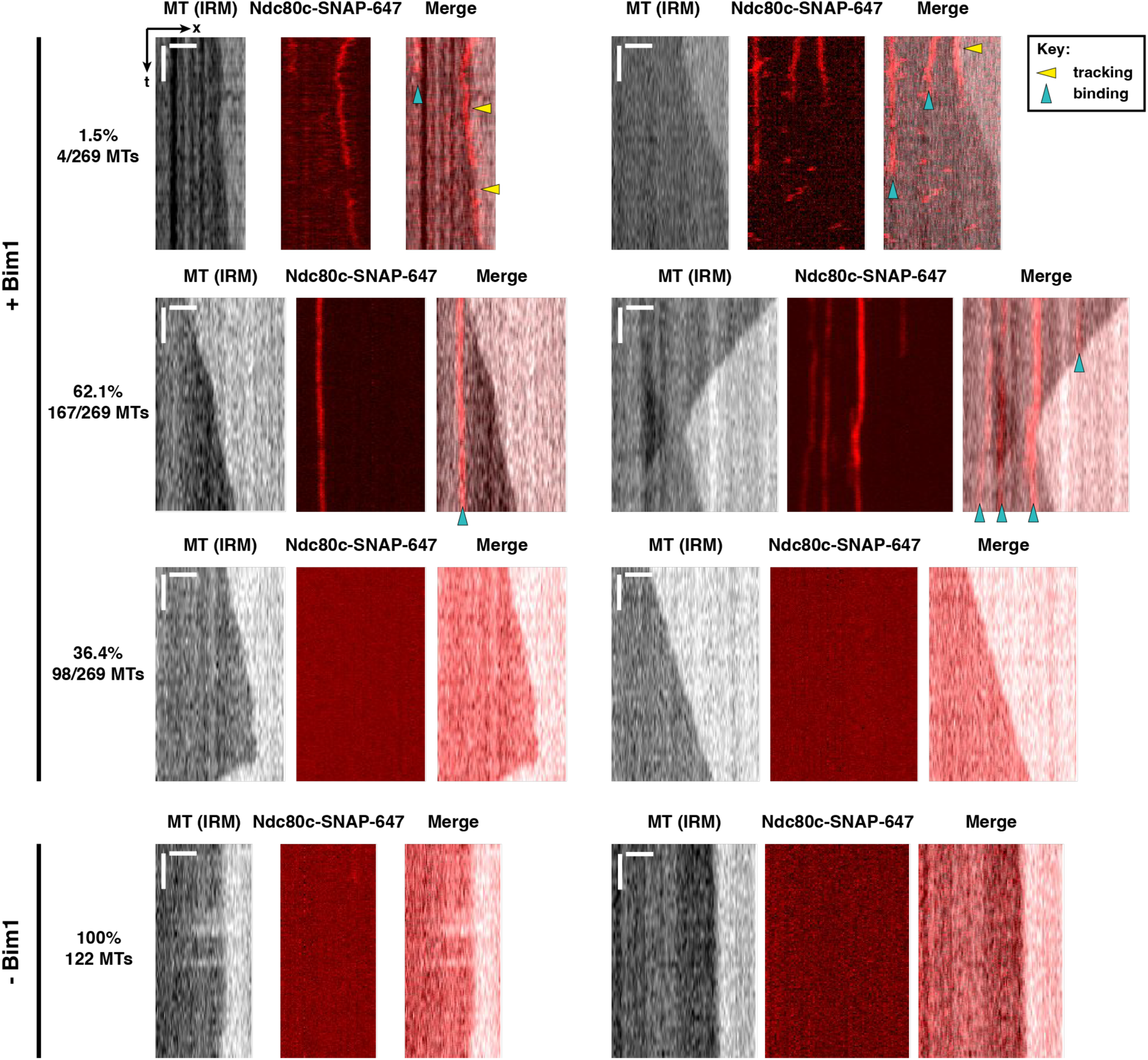
– *In vitro* reconstitution of Bim1–Ndc80c binding to dynamic microtubules. Representative kymographs showing dual IRM and TIRF imaging of Ndc80c-SNAP-647 binding to dynamic microtubules with (top) and without (bottom) unlabelled Bim1. 150 mM KCl was included in the imaging buffer to prevent binding to the microtubule lattice through Ndc80c. With Bim1, short-lived tip tracking (1.5%), lattice binding (62.1%) or no binding (36.4%) events were observed. No Ndc80c binding to microtubules was observed without Bim1. Scale bars 3 µm (x) and 30 s (t).

**Figure S8.**
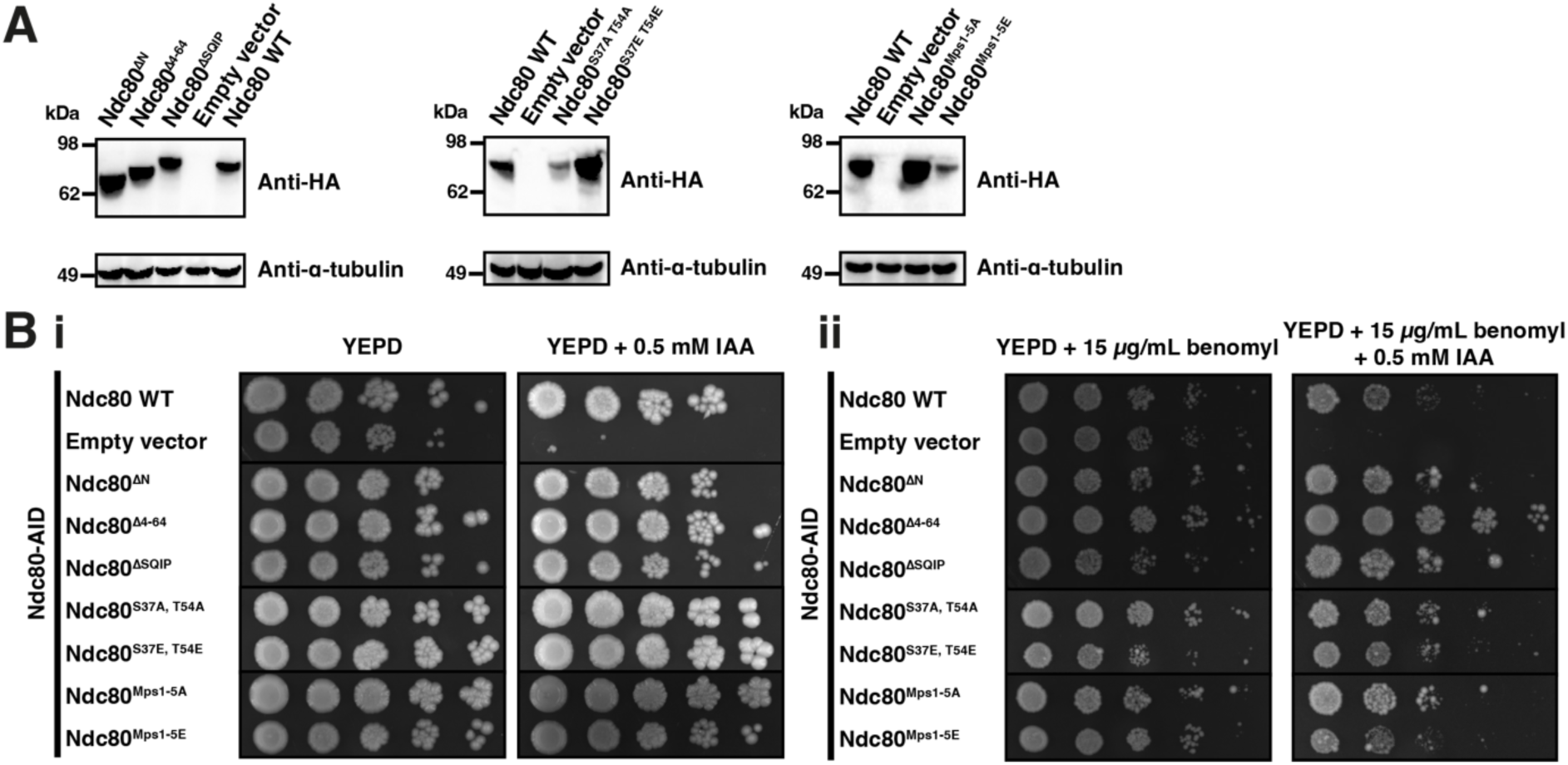
– *In vivo* investigations into the Bim1-Ndc80c interaction. **(A)** Detection of the indicated Ndc80-HA3 variants ectopically expressed in the Ndc80-AID strain (Muir et al., 2023) by Western Blot of whole cell extracts using an anti-HA (top) or anti-α-tubulin (loading control, bottom) antibody. **(B)** Auxin depletion assays for yeast strains expressing the indicated rescue alleles containing mutations in the Bim1-binding region of Ndc80 in the Ndc80-AID strain. Serial dilutions starting at OD 1.0 were spotted onto YEPD without (left) and with (right) auxin (IAA), and without (part i) or with (part ii) 15 µg/mL benomyl, and grown for 3 days at 30°C before imaging.

**Figure S9.**
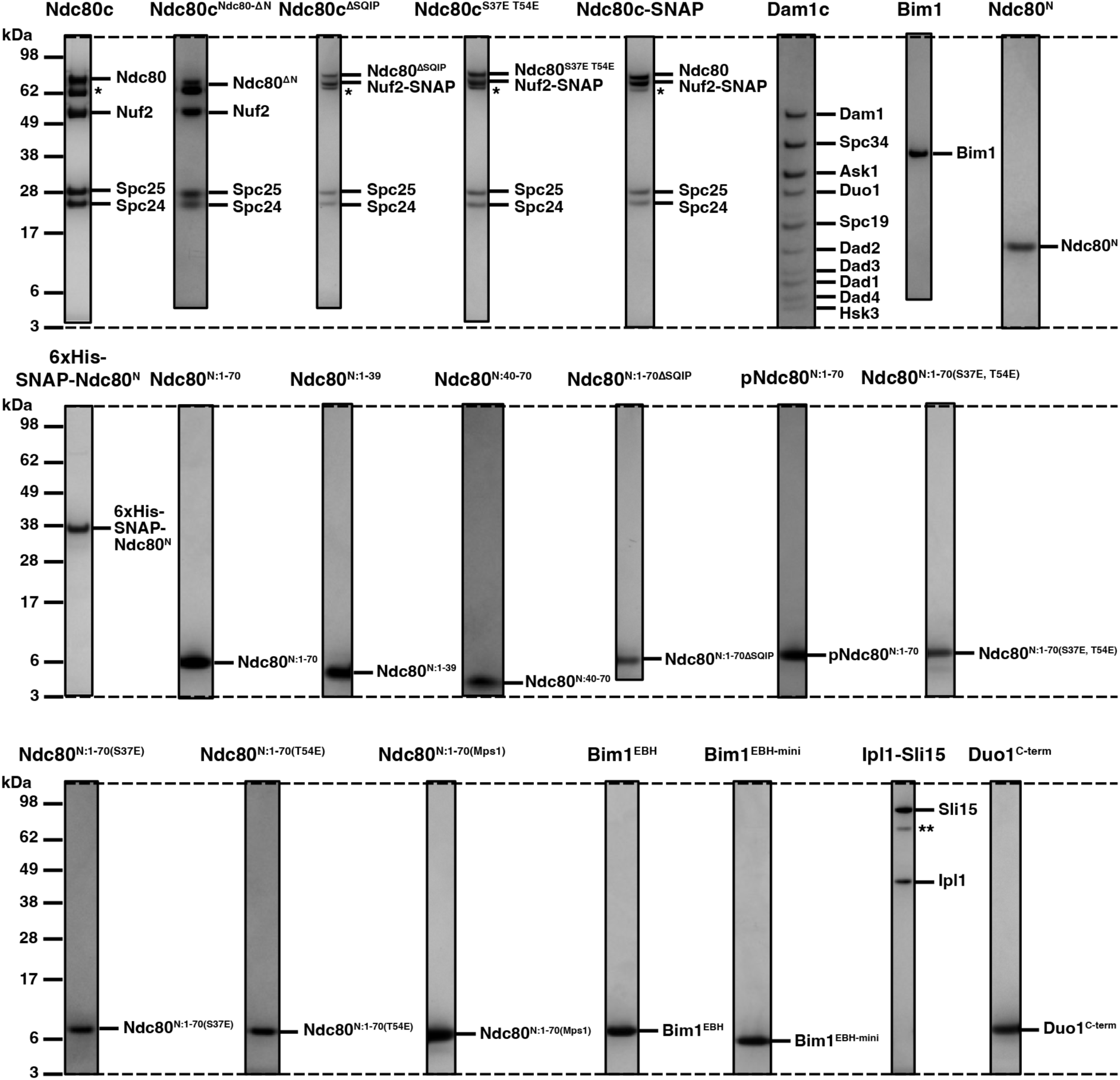
– Gallery of purified proteins. Representative SDS-PAGE gels from the final purification step for all purified proteins and complexes. Each row is aligned by the molecular weight ladder. All proteins were identified by mass spectrometry. * = Ndc80^ΔN^ (due to degradation of Ndc80^N^ during purification). ** = Sli15 degradation product.

**Table S1.** – Summary of ITC experiments performed in this study. All experiments tested for binding to Bim1^EBH^ except where specified. NB = no binding. ND = not determined.

| Sequence | K<sub>D</sub> (μM) | N |
| --- | --- | --- |
| Ndc80<sup>N</sup> | 4.2 | 1.0 |
| Ndc80<sup>N:15-29</sup> (SQIP peptide) | 178 | ND |
| Ndc80<sup>N:15-29ΔSQIP</sup> (mutant peptide) | NB | - |
| Ndc80<sup>N-helix</sup> (residues 39-54) | NB | - |
| Ndc80<sup>N:1-70</sup> | 4.0 | 0.95 |
| Ndc80<sup>N:1-70</sup> (with Bim1<sup>EBH-mini</sup>) | 7.8 | 1.01 |
| Ndc80<sup>N:1-70ΔSQIP</sup> | 41 | 0.97 |
| Ndc80<sup>N:1-39</sup> (primary binding site) | 21.5 | 1.39 |
| Ndc80<sup>N:40-70</sup> (secondary binding site) | NB | - |
| pNdc80<sup>N:1-70</sup> | 11.8 | 1.0 |
| Ndc80<sup>N:1-70(S37E, T54E)</sup> | 11.6 | 0.97 |
| Ndc80<sup>N:1-70(S37E)</sup> | 7.4 | 1.0 |
| Ndc80<sup>N:1-70(T54E)</sup> | 7.7 | 1.02 |
| Ndc80<sup>N:1-70(Mps1)</sup> | 142 | ND |
| Duo1<sup>C-term</sup> (residues 187-247) | 5.3 | 1.4 |

**Table S2.** – Yeast strains constructed for this study. ^▴^ denotes strains from Kyle Muir (Muir et al., 2023).

| Strain name | Genotype |
| --- | --- |
| Ndc80-AID<sup>Δ</sup> | *MATa ade2-1 his3-11,15 trp1-1 leu2-3,112 can1-100 ura3-1::ADH1-OsTIR1(pMK200, URA3), NDC80-mAID-FLAG<sub>3</sub>::G418* |
| Ndc80 WT<sup>Δ</sup> | *MATa ade2-1 his3-11,15 trp1-1 leu2-3,112 can1-100 ura3-1::ADH1-OsTIR1(pMK200, URA3), NDC80-mAID-FLAG<sub>3</sub>::G418, NDC80-HA<sub>3</sub>::TRP1* |
| Empty vector (TRP1)<sup>Δ</sup> | *MATa ade2-1 his3-11,15 trp1-1 leu2-3,112 can1-100 ura3-1::ADH1-OsTIR1(pMK200, URA3), NDC80-mAID-FLAG<sub>3</sub>::G418, pTRP1::TRP1* |
| Ndc80<sup>ΔN</sup> | *MATa ade2-1 his3-11,15 trp1-1 leu2-3,112 can1-100 ura3-1::ADH1-OsTIR1(pMK200, URA3), NDC80-mAID-FLAG<sub>3</sub>::G418, NDC80(Δ1-114)-HA<sub>3</sub>::TRP1* |
| Ndc80<sup>Δ4-64</sup> | *MATa ade2-1 his3-11,15 trp1-1 leu2-3,112 can1-100 ura3-1::ADH1-OsTIR1(pMK200, URA3), NDC80-mAID-FLAG<sub>3</sub>::G418, NDC80(Δ4-64)-HA<sub>3</sub>::TRP1* |
| Ndc80<sup>ΔSQIP</sup> | *MATa ade2-1 his3-11,15 trp1-1 leu2-3,112 can1-100 ura3-1::ADH1-OsTIR1(pMK200, URA3), NDC80-mAID-FLAG<sub>3</sub>::G418, NDC80(S22A, Q23S, I24A, P25S)-HA<sub>3</sub>::TRP1* |
| Ndc80<sup>S37A, T54A</sup> | *MATa ade2-1 his3-11,15 trp1-1 leu2-3,112 can1-100 ura3-1::ADH1-OsTIR1(pMK200, URA3), NDC80-mAID-FLAG<sub>3</sub>::G418, NDC80(S37A, T54A)-HA<sub>3</sub>::TRP1* |
| Ndc80<sup>S37E, T54E</sup> | *MATa ade2-1 his3-11,15 trp1-1 leu2-3,112 can1-100 ura3-1::ADH1-OsTIR1(pMK200, URA3), NDC80-mAID-FLAG<sub>3</sub>::G418, NDC80(S37E, T54E)-HA<sub>3</sub>::TRP1* |
| Ndc80<sup>Mps1-5A</sup> | *MATa ade2-1 his3-11,15 trp1-1 leu2-3,112 can1-100 ura3-1::ADH1-OsTIR1(pMK200, URA3), NDC80-mAID-FLAG<sub>3</sub>::G418, NDC80(T21A, S22A, S37A, T38A, T43A)-HA<sub>3</sub>::TRP1* |
| Ndc80<sup>Mps1-5E</sup> | *MATa ade2-1 his3-11,15 trp1-1 leu2-3,112 can1-100 ura3-1::ADH1-OsTIR1(pMK200, URA3), NDC80-mAID-FLAG<sub>3</sub>::G418, NDC80(T21E, S22E, S37E, T38E, T43E)-HA<sub>3</sub>::TRP1* |

